# Distributed encoding of action-mediated outcome drives consistent population dynamics during goal-directed reaching

**DOI:** 10.1101/2024.11.04.621878

**Authors:** Yangfan Peng, Öykü Okur, Carl Lindersson, Sasha Tinelli, Jeffrey Stedehouder, Rahul S Shah, Armin Lak, Charlotte J Stagg, Andrew Sharott

## Abstract

Anticipating the outcomes of actions is central to goal-directed behaviour, but how such expectations are encoded across the brain during ongoing movement remains unclear. To address this, we recorded spiking activity from cortical and subcortical regions using multiple Neuropixels probes simultaneously in head-fixed mice performing a water-reaching task. We found that distributed neural population dynamics were strongly modulated by the availability of reward beyond their encoding of forelimb kinematics. Principal component analysis revealed conserved population dynamics across brain regions and sessions that depended on reach amplitude and reward availability. Generalized linear models revealed outcome-related encoding within region-specific population dynamics, in addition to kinematic encoding, with the strongest outcome signals expressed in frontal cortico-thalamic regions. Unsupervised cluster analysis further identified outcome-encoding subpopulations that were enriched in frontal cortices and disproportionally contributed to the shared global latent dynamics. Together, these findings demonstrate that action-mediated outcome expectations are encoded in movement-related population dynamics that are shaped by functional clusters of neurons across the brain.

## Introduction

The brain generates movement to achieve specific goals. While the neural correlates of movements have been extensively studied in motor cortex, where neurons encode kinematic variables^1–5^, large-scale brain-wide recordings have shown that movement-related activity is distributed across widespread cortical and subcortical areas^6–9^. Similar widespread distributions have been reported for non-motor representations, such as sensory stimuli, choice and outcome^6,10–12^. These internal representations are often studied in temporally isolated task epochs outside the movement period, as it is challenging to dissect them from the strong and widespread activation during movement. Thus, it is unclear to what extent internally generated signals are expressed during ongoing action, and how they interact with motor dynamics at the single-neuron and population level. This question is especially relevant for goal-directed actions, where animals not only process sensorimotor feedback signals, but also must continuously adapt their internal representation of the goal based on the predicted and actual outcome of the action^13^.

Population level analysis can extract robust latent dynamics despite the diversity of single-neuron responses^14–17^. Such latent dynamics have been observed across brain regions and shown to reflect behaviourally relevant computations^18–21^, such as the representation of motor timing^22^. In motor cortex, distinct latent subspaces separate movement preparation, initiation, and execution^23–26^, while low-dimensional dynamics also correlate with spontaneous facial movements on the scale of seconds^27,28^. However, movement accounted for only 30-50% of variance in lower dimensions and higher dimensions appeared to be unrelated to motor output^27^, suggesting that non-motor signals contribute significantly to population dynamics during movement. This raises the question of what internal representations underlie these dynamics during ongoing movement at millisecond timescale, and how such global population-level activity relates to the functional tuning of individual neurons within different brain regions^29^.

One potential internal variable shaping ongoing population dynamics during fast goal-directed movements is the anticipation of reward or outcome, as seen in dopamine signals that track reward proximity during locomotion^30^. While outcome-related activity has been well established in classical reward-processing regions such as striatum, prefrontal cortex, and midbrain^31,32^, recent studies have shown outcome-related signals distributed across the brain, including motor cortex and subcortical regions^8,33–35^. However, most of these studies have focused on reward expectation prior to movement^36,37^, or on outcome encoding at the time of reward delivery or during reward consumption^8,38^. These later periods can be confounded by consummatory behaviours such as licking. In addition, outcome expectation can act as an “actionable representation”, a dynamic signal that predicts and guides the ongoing movement^39^ , yet this has not been systematically studied across multiple brain regions during the action itself. Thus, it remains unclear how outcome is represented during the ongoing execution of goal-directed actions.

To address these questions, we recorded the simultaneous spiking activity of over a thousand neurons across motor and non-motor brain regions during a reaching task in head-fixed mice. We found task-related stereotypical population activity in a latent subspace that was preserved across regions and animals. Using unsupervised clustering and generalized linear modelling, we further identified functional subpopulations modulated by reward availability during movement, independent of reach kinematics, with strong enrichment in frontal thalamocortical regions. Our findings show that functional subpopulations contribute to population dynamics within and across regions that encode outcome-related representations during movement.

## Results

### To study movement-related neuronal population dynamics, we performed high-density Neuropixels recordings in head-fixed mice performing a forelimb reaching task

To initiate a trial, the animal had to rest its left paw on a metal bar for a holding period of 3-6 seconds to trigger the presentation of a 4-5 µl sucrose water droplet which was signalled by an 80 ms long cue tone (Fig. 1a, task adapted from previous study^40^). The animal then immediately reached towards the droplet, grabbed it, and brought the droplet to the mouth (Fig. 1a). Using electrical detectors (pyControl^41^) and automated pose estimation in 3D space (DeepLabCut^42^ with Argus3D^43^), we were able to extract the times of reach onset (bar release), reward retrieval (droplet disappears) and the movement trajectory of the forelimb joints at high temporal and spatial resolution (Fig. 1b, Fig. S1, Suppl. Video 1). After training (see Methods), the animal learned to reach up to 200 times upon cue within a 30-minute recording session. Cued reaches were highly consistent across trials, sessions, and animals with similar reaction times (234 ms [170-390 ms], median [interquartile range]), movement times (bar to spout: 133 ms [83-211 ms]) and similar movement trajectories (Fig. S1). In total, we recorded 5738 cued reaches across 38 sessions (151 ± 59 trials per session, mean ± SD) from 7 animals.

**Figure 1:**
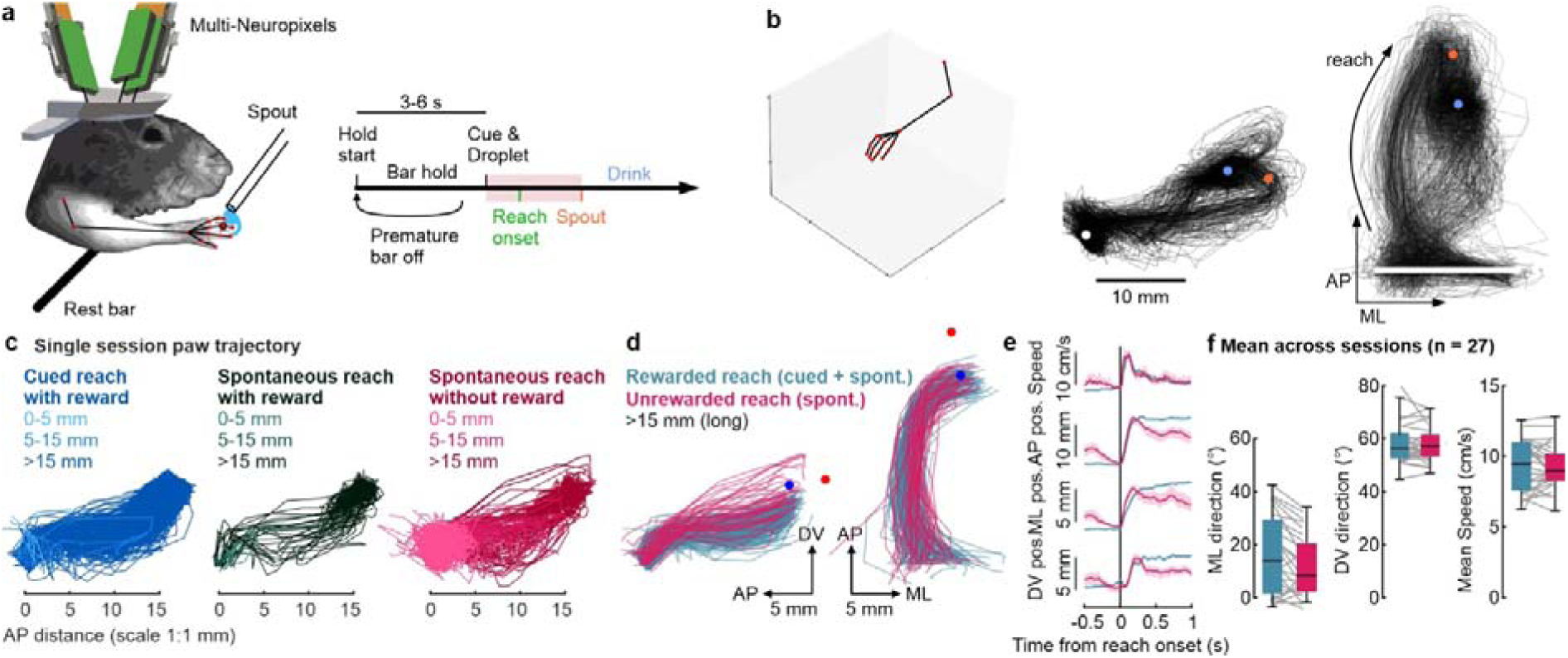
Performance on a goal directed reaching task leads to cued and spontaneous movements with different reward-related outcomes. **(a)** Schematic of head-fixed recording setup and water reaching task. Cue and droplet are presented after 3-6 s holding period, red shaded area illustrates reaction time until reach onset (bar release) and reach duration until spout touch. **(b)** Front and side video allowed projection of automatically estimated joint position into 3D space. Black trajectories represent movement of left digit II from the side view and top view during one session. White dot and line represent rest bar. Orange dot shows spout position, blue dot indicates location of reward consumption. **(c)** Side view of movement trajectories of left digit II from example session at 1:1 scale. Holding bar is located at bottom left, droplet position is at top right. Colors indicate different movement types, lightness categorizes maximum reach distance within 500 ms. **(d)** Example session data showing top and side view of movement trajectories of long reaches during the “approach period”. Reaches were color coded by the presence (teal) or absence of reward (pink), irrespective of cue. **(e)** Example session trial-averaged traces of reaches in both conditions for kinematic parameters. **(f)** Boxplots show distribution of session-averages (gray lines) in kinematic features between long rewarded (teal) and long unrewarded reaches (pink). ***p < 0.001, calculated by LME.

In addition to cued reaches, mice performed spontaneous reaches from the bar that occurred later than 1 s after the cue (n = 7,245, Fig. 1c). While cued reaches were always executed in the presence of reward, spontaneous reaches were performed either while the droplet was still present (19% “with reward”) or already absent because of prior retrieval (81% “without reward”). We further found that the mice performed reaches of varying reach distances, sometimes just briefly lifting the paw and immediately returning to the bar (< 5 mm, short) while more often performing a full reach that allows retrieval of the droplet (> 15 mm, long) (Fig. 1c). As we primarily study outcome-related activity, we focused our subsequent analyses on long reaches, comparing rewarded reaches, including cued and spontaneous reaches, with unrewarded spontaneous reaches. Movement trajectories of these long reach types were highly consistent (Fig. 1d-e). We compared the reach trajectories across sessions and found no significant differences (p > 0.05 in LME) in rewarded versus unrewarded reaches with respect to speed (9.3 ± 1.8 vs. 9.3 ± 1.6 cm/s, mean ± SD) and reach directions (ML direction: 17 ± 15 vs. 12 ± 12°; DV direction: 58 ± 8 vs. 58 ± 7°). The only significant difference was a small 4% reduction in trajectory length (25 ± 3 vs. 23 ± 2 mm, p < 0.001; Fig. 1f). Furthermore, mice also performed grooming movements which we could isolate using the 3D movement trajectories (see Methods). This diversity of movement types under different conditions enabled us to disentangle neural representations of outcome expectation from movement execution.

### We found positively and negatively modulated neurons across all recorded regions

To simultaneously record neuron spiking activity, we inserted two to three Neuropixels probes into the brain (probe-1: premotor cortex corresponding to rostral forelimb area, medial prefrontal cortex, orbitofrontal cortex, olfactory areas; probe-2: primary motor cortex corresponding to caudal forelimb area, dorsolateral striatum; probe-3 in mice 5-7: hippocampus, thalamic nuclei, hypothalamus; Fig. 2b). After spike sorting, quality control and region assignment (Fig. S2, see Methods), we obtained a pooled dataset of 22,036 neurons exhibiting a wide range of activity patterns during cued reaches (Fig. 2c). We included a mean of 1,574 neurons per region (range: 326 to 4,129, Fig. 2f). To broadly characterize their task-related modulation, we first focused on cued reaches and determined the relative change in trial-averaged firing rate 500 ms after the cue relative to 250 ms baseline before the cue (modulation index, see Methods). While we observed highly heterogenous and skewed distributions of modulation across all recorded brain regions (Fig. 2d-e, Fig. S3), region-specific differences were evident when analysed at the session-level (Fig. 2f). Sensory, interlaminar and mediodorsal thalamus showed the highest fraction of positively modulated neurons (∼ 50%), with lower fractions in non-thalamic cortical and subcortical regions. This effect mostly arises from neurons with intermediate firing rates (1–10 Hz, Fig. S3). Negatively modulated neurons were found across all recorded regions. While the fraction of negatively modulated neurons was lower than that of positively modulated neurons in most regions, the hippocampus exhibited the strongest negative modulation, a pattern that was specific to sparsely firing neurons (<1 Hz, Fig. S3).

**Figure 2:**
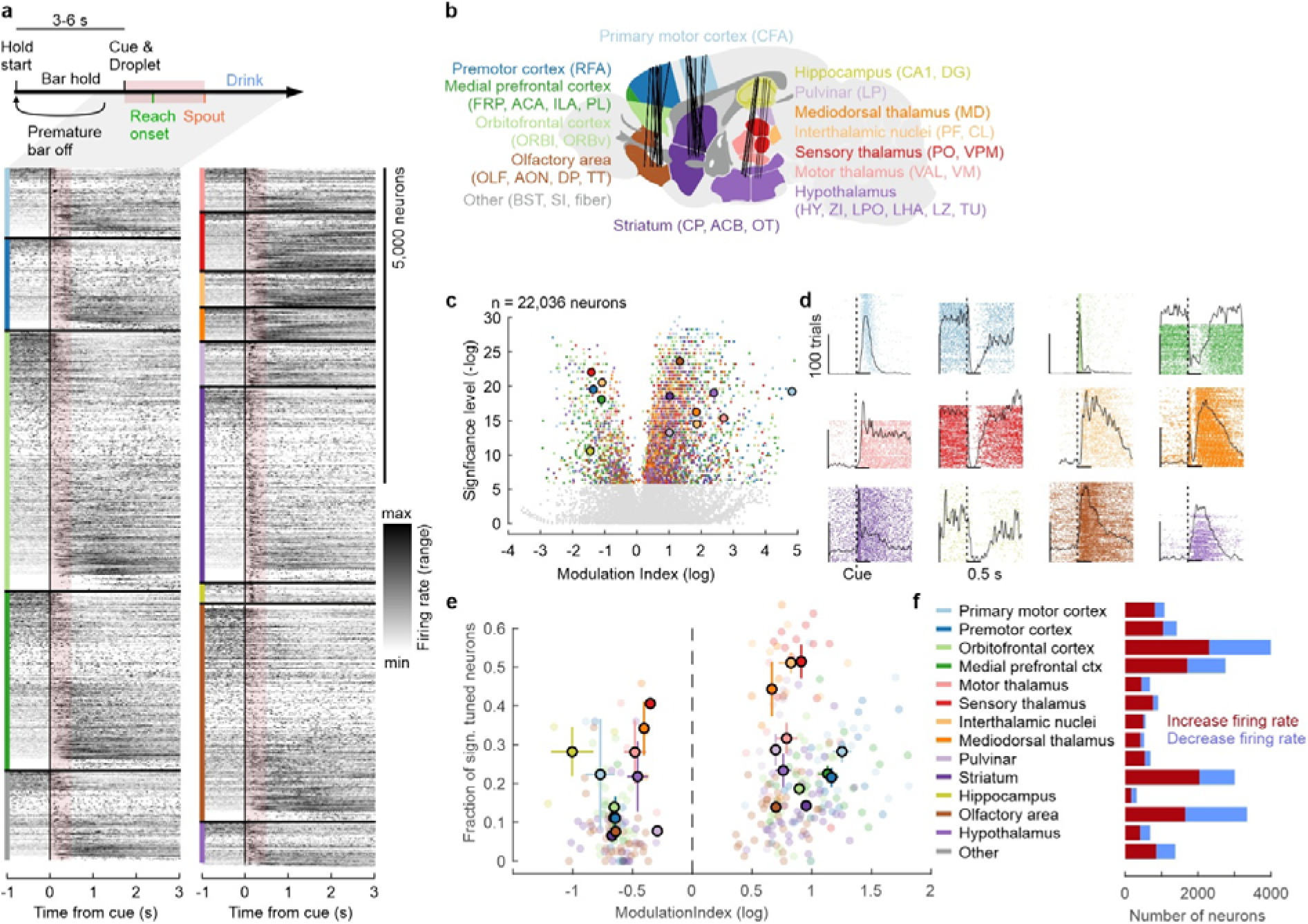
Heterogeneous modulation of neuron activity across regions during reaching. **(a)** Peri-spike timing histogram (PSTH) shows trial-averaged and range-normalized firing rate of all neurons (rows) from all recording sessions, aligned to time of cue and sorted by brain regions and product of modulation index and significance level. Shaded red interval indicates 500 ms window used to compute modulation index. **(b)** Schematic of sagittal brain atlas showing recorded region categories in colour. Abbreviation in parentheses indicate included subregions according to Allen Mouse Common Coordinate Framework. Black lines represent Neuropixels trajectories from different sessions approximating positions of active electrodes. **(c)** Volcano plot shows cue modulation index (log) and significance level (-log10) for each recorded neuron, color-coded by its brain region. Grey neurons are not significant after Bonferroni correction (p > 1.3 x 10^-06^). Circled dots indicate example neurons from each region with spike rasters shown on the right (matching colours). **(d)** Spike rasters show spiking activity (dots) across all cued reach trials (rows) from individual neurons, aligned to time of cue (dashed vertical line). Vertical scale bar: 100 trials, horizontal scale bar: 1 s. **(e)** Scatter plot shows fraction of significantly positively modulated (index > 1) and negatively modulated (index < 1) neurons in each region (colour). Each transparent dot indicates an individual recording session, circled dots represent mean and standard error of mean across sessions per region. **(f)** Bar graph shows pooled number of neurons recorded in each region, subdivided into their activity change upon cue.

### To better interpret the heterogeneity of single neuron responses, we next analysed the neuronal population activity using a dynamical systems framework^18^

Previous studies have shown that movement-related latent population dynamics can exhibit stable patterns despite heterogeneity of individual neuronal activity, reflecting the capability of the neural network to maintain functional coherence despite diverse cellular responses^19,23,44,45^. To identify the most dominant and stable neuronal population trajectories in our data, we performed principal component analysis (PCA)^17^ on the activity of simultaneously recorded neurons of all regions from each session (Fig. 3a, Fig. S4a-d). We first focused only on cued long reach trials and aligned them to reach onset (bar release). The PCA was performed on the trial-average of normalized firing rate (z-score across entire session) around reach onset (−1.5 s to 3 s) from all neurons of each session, using 20/80 cross-validation (PCA fit on training trials and evaluated on held-out test trials; see Methods). The first principal component (PC1) explained 22 ± 5% of the variance (Fig. S4e) and increased before reach onset while peaking after it (Fig. 3a). The second principal component (PC2) explained 14 ± 3% of the variance and exhibited a peak around reach onset. The decay in explained variance (or eigenvalues) across components followed a power law decay with an exponent of 1.10 [1.09 to 1.10] (Fig. S4e).

**Figure 3:**
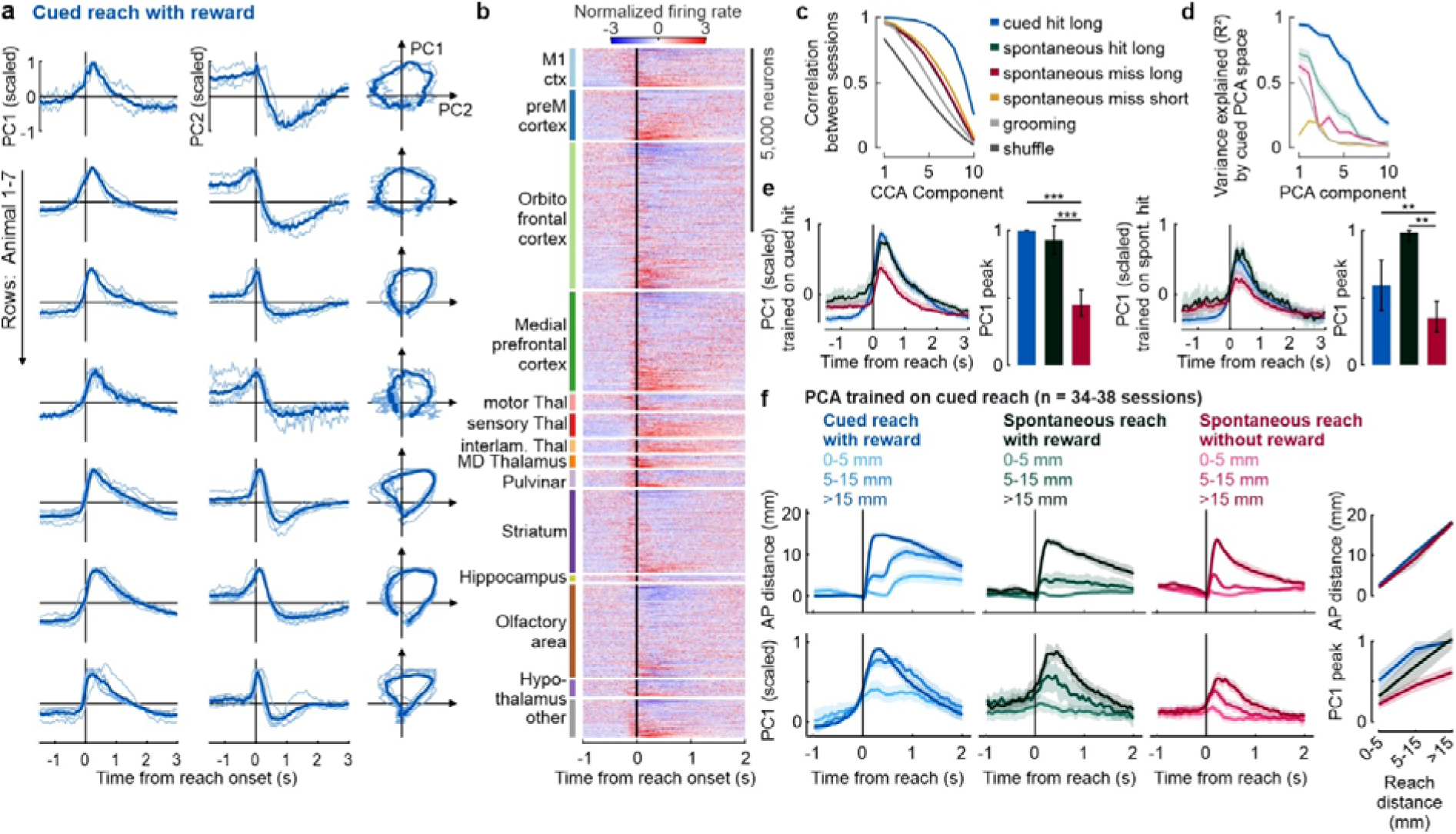
Consistent reach-related latent dynamics across sessions and regions are modulated by reward availability and reach distance. **(a)** Line plots show trial-averaged PC1 and PC2 dynamics during cued long reaches of each session (light blue lines) and animal-wise average (dark blue line) across animals (rows). PC dynamics were obtained from a PCA computed on trial-averaged activity using 20/80 cross-validation and scaled to the maximum peak of each session. The rightmost column shows state-space trajectories of individual sessions and of the animal-wise average. Three sessions with cross-validated R² < 0.8 were excluded. Shaded areas represent bootstrapped 95% confidence intervals (CI). **(b)** PSTH shows trial-averaged and z-scored firing rate of pooled neurons (rows) during cued reaches from all sessions, sorted by PC1 coefficient and brain region. **(c)** CCA was performed between each pair of sessions to quantify degree of similarity between sessions after alignment, lines indicate average canonical correlation across all session pairs for the top 10 CCA components of PCA dynamics calculated on trial-averaged activity of different movement types (colors) or shuffled control (dark gray). **(d)** Line plots show the mean fraction of variance explained (R²) by the top ten PCs across sessions for a PCA trained on cued long reaches. Thin lines indicate cross-condition projections, showing how well activity from other movement types (colors) is captured in this PCA subspace. Thick line indicates within-condition projections (cued long reaches), where R² reflects the variance explained for held-out test trials (20%) projected into a PCA subspace computed from training trials (80%, 20/80 cross-validation). **(e)** Trial-averaged PC1 traces of different reach types (color code as in panel c) were obtained by projecting activity into the PCA subspace trained on either rewarded cued long reach (left) or rewarded spontaneous long reach (right) and normalized within each session to the peak amplitude of PC1 in that training subspace. Thick lines represent across-session mean. As in panel d, within-type projections show 20/80 cross-validation. Bar plots show average of normalized PC1 peak amplitude across sessions, error bars indicate 95% CI. Asterisks show significant differences based on sign-rank test: **p < 0.01, ***p < 0.001. **(f)** Top: Movement position along the anterior-posterior (AP) axis for different movement types (colors), aligned to movement onset. Lines indicate the mean across sessions (lightness show reach distances). Bottom: Corresponding activity along PC1 trained on cued long reaches, normalized to the peak amplitude of long cued reaches within each session. Only sessions with at least five trials per reach type were included. Shaded areas represent 95% CI.

### The latent dynamics are consistent across sessions and animals

To assess the consistency of the population dynamics across sessions, we computed pairwise alignment of the top ten PCs from different recording sessions using both canonical cross correlation analysis (CCA) and Procrustes distance (see Methods)^46,47^. These are well-established approaches to quantify the similarity of neural latent dynamics across different sessions^47–49^. As a baseline control, we applied the same alignment analyses to shuffled data (n = 100 per session), where PCA was performed on the population firing rate in randomly selected time windows of equal length (4.5 s). This control sets a lower bound for expected similarity that arise from task-independent neuron-to-neuron covariance^48^. Latent dynamics were highly consistent across sessions, both within and across animals (Fig. 3a-b). CCA showed highly correlated dynamics within the top canonical components (CC) across sessions (Fig. 3c) and the Procrustes distance in the data was significantly lower compared to control (d_within_animal_ = 0.12 ± 0.12 and d_across_animal_ = 0.17 ± 0.11 vs. d_control_ = 0.83 ± 0.03; p < 0.001). These results suggest that population activity across different sessions and animals are dominated by preserved latent dynamics even though each session recorded from putatively different neurons.

### The latent dynamics are strongly dependent on outcome and movement kinematics

To assess the generalizability of latent subspaces across different reach types, we projected trial-averaged activity from each reach type into PCA subspaces trained on other reach types and quantified the variance explained (Fig. 3d, Fig. S4g). PC1 trained on cued long reaches explained most of the variance both in cued (94%, 20/80 cross-validated) and in spontaneous long reaches performed with reward (72%), but less variance in spontaneous long reaches without reward (63%, Fig. 3d). Consistent with this, when projecting activity into the latent subspace trained on cued long reaches, the peak amplitude of PC1 was comparable between rewarded (hit) cued and spontaneous reaches. In contrast, spontaneous reaches executed in the absence of reward (miss) exhibited a significantly reduced PC1 peak (Fig. 3e). Furthermore, the PC1 peak of rewarded reaches, both cued and spontaneous, was significantly higher than unrewarded reaches in the latent subspace trained on rewarded spontaneous long reaches (Fig. 3e). These results indicate that the dominant latent subspace generalizes across rewarded movements and is strongly dependent on the successful outcome.

In addition to outcome-related modulation, we found that PC1 peak amplitude scaled with reach distance, increasing from short to intermediate to long reaches across all reach types (Fig. 3f). Importantly, outcome-related modulation was present across reach distances and explained 26% of PC1 peak variability in a linear mixed effect model that took movement distance, cue and outcome into account (see Methods). Thus, while movement-related activity in dominant latent dynamics has been established across multiple brain regions^6,7,27^, our results indicate that population activity is strongly modulated by outcome beyond pure movement kinematics which we disentangle in the following analyses.

### Outcome and kinematics uniquely contribute to population dynamics

Our previous analyses on the trial level showed that the dominant population activity was dependent on outcome during ongoing movement, suggesting an outcome representation that is distinct from the activity related to reward consumption. However, as the animal is continuously executing goal-directed reaching movements during the period of reward presentation until the droplet is consumed, movement kinematics and outcome expectation are strongly correlated. To disentangle these factors, we leveraged the timing variability of reward consumption, the kinematic variability of reach movements and the fact that they were performed in the presence and absence of reward to fit a generalized linear model (GLM). The model predictors included continuous kinematic variables (bar distance, speed), kernels for events (cue, movement onset) and kernels for time periods of reward presence and consumption (Fig. 4a). For each predictor we computed its share of explained variance (R² contribution) of the full model R² on the neural activity that is uniquely attributable to that variable calculated using a leave-one-out approach^50^ (see Methods). Predictors showed minimal multicollinearity (VIF range across conditions: 1.00-1.19), meaning that kinematic and outcome-related predictors were sufficiently separable and that comparison of relative contributions from partial models is possible. This allowed us to isolate the contribution of specific outcome variables to neural activity after regressing out movement kinematic parameters.

**Figure 4:**
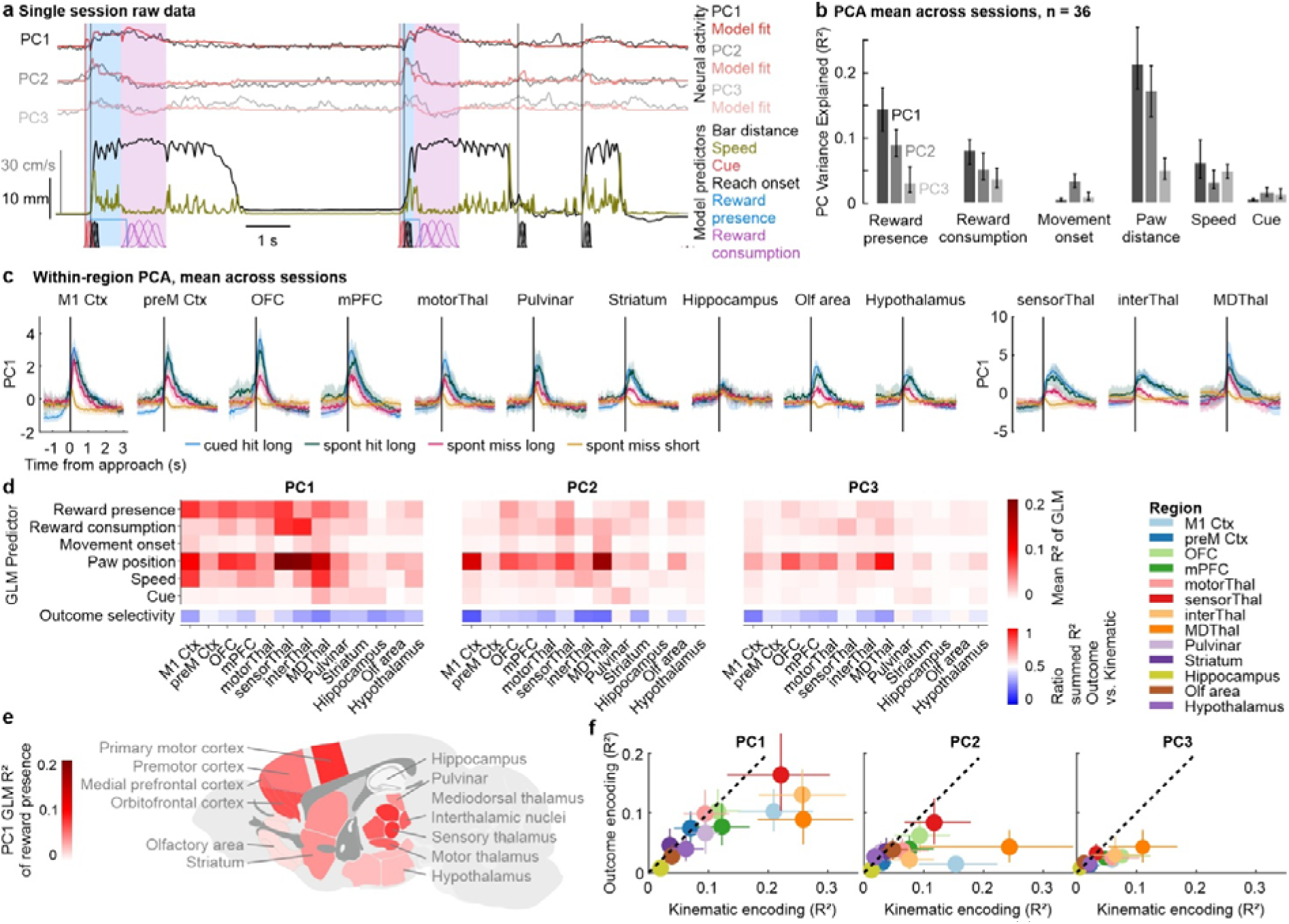
Independent contribution of outcome and kinematic encoding to population activity across regions. **(a)** Example continuous data of one session with PC1-3 expression over time (trained on cued reaches, gray lines) overlaid with GLM fits (red lines) and behavioural predictors below. Lines show continuous predictors, including position and speed. Vertical lines indicate event times, such as cue and reach onset and shaded bars show time periods of reward presence and consumption. These discrete events are modelled as kernels or step functions. **(b)** Bar graphs show mean R² (unique contribution based on leave-one-out approach, see Methods) per predictor for PC1-3 across sessions (n = 36 sessions, error bars indicate bootstrapped 95% CI). **(c)** Trial-averaged region-specific PC1 traces of different reach types (colors), obtained by projecting activity into the PCA subspace trained on within-region activity during rewarded cued long reaches. Lines represent across-session mean, shades represent 95% CI. **(d)** Heatmap shows session-mean of unique R² for each predictor (rows) in GLM fitted on PC1-3 activity calculated within individual regions (columns). The bottom row shows the outcome selectivity, defined as the ratio of explained variance by outcome predictors (reward presence and consumption) relative to kinematic predictors (movement onset, position, speed) for each region. **(e)** Sagittal schematic of recorded brain regions, color-coded by the across-session mean R² of reward presence on region-specific PC1 activity. **(f)** Scatter plots show relative amount of variance explained by kinematic vs. outcome encoding parameters on top PCA components within each region (colors). Dots represents mean across sessions, error bars show 95% CI, dashed line represents slope of 1 with equal contributions.

We first fitted the GLM on the continuous expression of PCA components that were calculated based on trial-averaged activity of all recorded neurons during cued long reaches. Across sessions, the GLM showed that paw distance to bar was the dominant predictor of PC1 activity (R² = 0.21 [0.17 to 0,27], mean [95% confidence interval], Fig. 4b). Crucially, reward presence was the second-strongest predictor with a unique contribution of R² = 0.14 [0.11 to 0.18]. PC2 showed slightly lower contributions of both kinematic and outcome predictors but exhibited the highest contribution of the movement onset kernel compared to the other top components. PC3 showed the lowest contribution of paw distance and outcome parameters, but a relatively high contribution by movement speed. These results confirm that the dominant global latent dynamics reflect a combination of both movement kinematics and action outcome, with each contributing uniquely to the population activity.

### Frontal corticothalamic regions exhibit strongest outcome encoding

All analyses so far focused on the latent dynamics extracted from pooling neuronal activity across all recorded regions. We next sought to estimate to what extent the global PC1 reflects local population dynamics within each region. For this, we performed PCA separately on simultaneously recorded neurons from individual regions (local PCA) and found that local PC1 coefficients of each neuron strongly correlated with the corresponding global PC1 coefficient (Fig. S5b-c). Consequently, projected neurons from one region onto the global or local PC1 dimension showed largely consistent dynamics across regions (Fig. 4c, Fig. S5a). To increase interpretability, we focused on the region-specific latent population activity and analysed their outcome and movement encoding. We found that region-specific PC1 activity was also strongly dependent on outcome and kinematics. All regions except for hippocampus exhibited higher PC1 peak in long vs. short reaches (Fig. 4c). Outcome modulation, as shown by higher PC1 peak in rewarded vs. unrewarded long reaches, was most prominent in prefrontal cortices (preM, OFC, mPFC, M1), thalamic areas (sensory, interthalamic), Striatum, Olfactory area, and Hypothalamus. We next fitted the above-described GLM to the continuous expression of these region-specific latent dynamics and found that outcome was uniquely encoded in PC1 dynamics across all regions except for the hippocampus (Fig. 4d, 4e, see Methods). At the same time, movement kinematics were also uniquely encoded in the region-specific PC1 with the strongest encoding in primary motor cortex and specific thalamic areas (sensory, interthalamic, mediodorsal). Most regions exhibited slightly more kinematic than outcome encoding, while premotor cortex and motor thalamus showed a balanced encoding of kinematic and outcome (Fig. 4f). Higher dimensions exhibited weaker encoding of outcome, but a relatively high encoding of kinematics, most of which relating to paw position (Fig. 4d, 4f). These results confirm that the dominant PCA latent dynamics within almost all regions (except hippocampus) jointly encode outcome and kinematics with a preference towards outcome encoding in frontal corticothalamic regions.

### Outcome representation persists in the pre-reward movement window even when movement trajectories are matched

To further establish that outcome-related activity is present during goal-directed movement before reward consumption, and that it is not driven by subtle kinematic differences, we focused on the “approach period”, defined as the period of continuous decrease in paw-to-droplet distance (Fig. 5a). As mentioned above, we found no significant differences between rewarded versus unrewarded reaches with respect to speed and reach direction on a session level (Fig. 1f, p > 0.05 in LME). Despite these similarities between reach types, individual reaches exhibited substantial within-session variability in trajectory length (SD = 3 mm, mean standard deviation across sessions), mean speed (SD = 2.8 cm/s), ML angle (SD = 26°), and DV angle (SD = 14°, Fig. 5b). This within-session variability enabled us to perform a trial-by-trial comparison of outcome-related activity between reach trajectories with varying degrees of similarity. To this end, we computed the difference in average global PC1 activity during the approach period between all possible rewarded–unrewarded long-reach pairs within each session and grouped these trial pairs by similarity in kinematic features. Across all kinematic features, PC1 activity was consistently higher for rewarded than unrewarded reaches (Fig. 5c). Importantly, this outcome-dependent activity difference persisted even for reach pairs with high kinematic similarity (Δkinematic feature ≈ 0) and was significantly greater than in shuffled controls (Fig. 5c). Taken together, these results indicate that outcome encoding in latent population dynamics reflects an internal representation during ongoing movement that cannot be explained by differences in movement kinematics.

**Figure 5:**
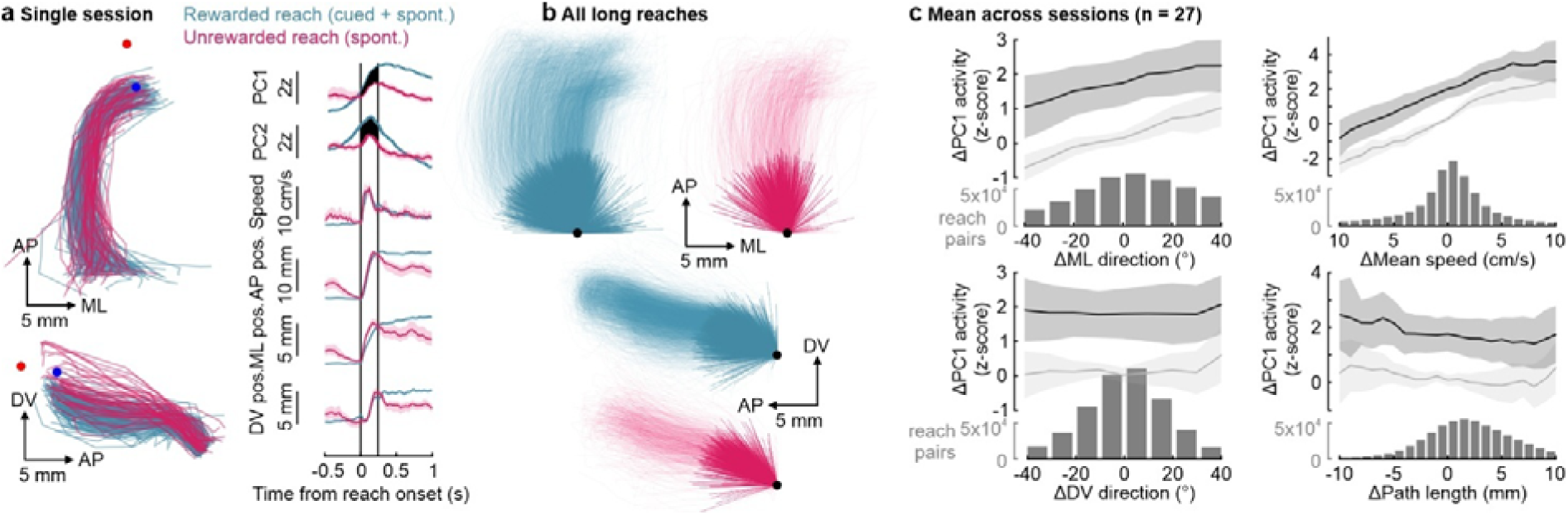
Kinematic similarity analysis shows persistent outcome encoding in matched reach trajectories. **(a)** Example session data showing top and side view of movement trajectories of long reaches during the “approach period”, defined as a period of continuous decrease in paw-droplet distance. Reaches were color coded by the presence (teal) or absence of reward (pink), irrespective of cue. Traces show trial-averaged data from this example session of reaches in both conditions for z-scored PC1-2 activity, speed and position across all axes. Vertical black lines show the approximate boundaries of the “approach period” during which the averaged PC Δfiring rate was calculated (black area between curves). Note that the approach period was individually determined for each reach pair (see Methods). Shaded error indicates bootstrapped 95% CIs. **(b)** Overlay of all rewarded (teal) and unrewarded (pink) long reach trajectories during the approach period, shown in top and side view and aligned to start position (black dot). Data include sessions with at least five trials of each reach type (n = 27). Initial reach angles (thick lines) were calculated from start point to the point where the forepaw had travelled 10 mm along the path. **(c)** Within each session, rewarded-unrewarded reach pairs were binned by difference in kinematic features (x-axis of each panel) and the mean difference in PC1 activity (y-axis) was calculated for each bin. Bins close to zero on the x-axis represent reach pairs had nearly identical values in the respective kinematic feature. The black lines show the across-session difference in PC1 calculated from individual session-means, shaded error represents 95% CI at session level. Light gray lines show the across-session mean difference after shuffling reward labels. Histograms below each line plot show total number of reach pairs in each bin from all included sessions (n = 27).

### A distinct subpopulation of neurons encodes outcome-related information during movement

While PCA showed that outcome and kinematics are jointly encoded at the population-level, it does not reveal whether specific subpopulations of neurons contribute to the different behavioural variables. To identify functionally interpretable subpopulations, we applied an unsupervised clustering method (Rastermap) to the continuous firing rate of all neurons recorded during a session^51^. This allowed us to cluster neurons based on shared temporal firing patterns across the entire session, independent of trial structure (Fig. 6a, see Methods). We found that the average activity of one cluster of neurons was highly correlated to the onset of reward consumption (Fig. 6a). The average firing rate of this cluster sharply increased upon movement onset and peaked at the time when the paw reached the mouth. In rewarded reaches, this time point corresponds to the start of reward consumption. Notably, this peak firing rate was similar between cued and spontaneous rewarded reaches and markedly reduced in unrewarded long reaches. This outcome-dependent difference in trial-averaged activity was consistently observed across sessions, when selecting for the most outcome-encoding cluster, which we describe below (Fig. 6b). To identify these outcome encoding clusters in each session, we applied the previous GLM approach to each Rastermap cluster to determine which behavioural variables best explained their activity (Fig. 6c, 6d). For each session, we labelled the cluster with the highest R^2^ contribution from the reward presence predictor as the top reward presence cluster (RP1). Across sessions, RP1 clusters explained 0.15 [0.11 to 0.18] (mean [95% confidence interval]) of the variance related to reward presence (Fig. 6d). Movement kinematics were also encoded, but at a lower level (paw distance: 0.1 [0.07 to 0.17], speed: 0.07 [0.05 to 0.09]). Contributions from other variables, including reward consumption, were smaller (R² < 0.05). Thus, combining unsupervised clustering of neurons with GLM analysis revealed a distinct neuronal subpopulation that exhibited outcome encoding during ongoing movement that could not be explained by differences in kinematics alone.

**Figure 6:**
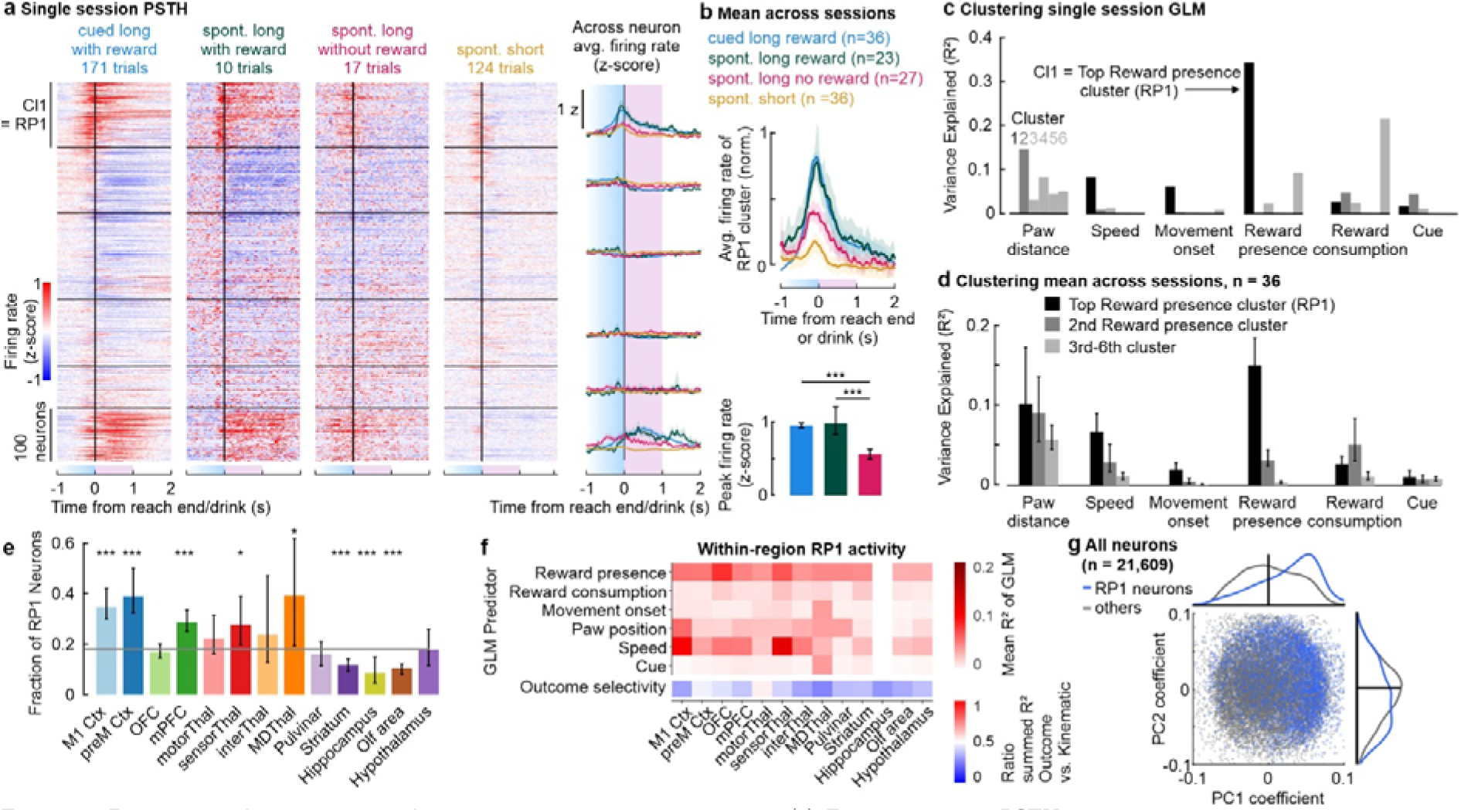
Region-specific prevalence of outcome-encoding subpopulation. **(a)** Example session PSTH showing trial-averaged and normalized firing rate of neurons sorted and clustered by Rastermap (horizontal lines, see Methods). Activity during different reach types are shown (columns). Neural activity was aligned to the end of reach, defined as the end of the “approach” period. In rewarded reaches, this time point corresponds to the time when the droplet is brought to the mouth, separating the prior time period of reward presence (blue shaded area) from the subsequent period of reward consumption (purple). Traces on the right show overlaid trial-averaged activity of each cluster during different reach types (color). (b) Traces show the across-session mean activity of the top reward-presence cluster (RP1), defined in each session as the cluster with the highest GLM encoding of reward presence (see panel c and Methods). RP1 activity during different reach types (colors) are overlaid. Bar plots below show across-session mean of RP1 peak firing rate across long reach types. Shaded areas and error bars represent 95% CI. Asterisks show significant differences based on sign-rank test: ***p < 0.001. **(c)** Bar graphs show explained variance (R²) of each GLM predictor rastermap clusters 1-6 (adjacent bars) in the example session from panel a. The cluster with the highest reward presence R² (cluster 1) is labelled as “Reward-Presence 1” (RP1, black bars) **(d)** Bar graphs show mean R² per predictor for RP1 (black) and other clusters that were sorted by the amount of variance explained by reward presence within each session (n = 36 sessions, error bars indicate bootstrapped 95% CI). **(e)** Bar plot shows across-session mean RP1 fraction relative to all neurons within each region (x-axis, color) and session. Horizontal gray line represents the average overall fraction of RP1 neurons (18%). Error bars indicate bootstrapped 95% CI. Statistical significance was determined using an LME with animal and session as nested random effect, *p < 0.05, ***p < 0.001. **(f)** Heatmap shows session-mean of unique R² for each predictor (rows) in GLM fitted on average RP1 activity within each region (columns). Outcome selectivity represents the ratio of explained variance between outcome to kinematic encoding. **(g)** Scatter plot of PC1 and PC2 coefficients of RP1 (blue) and other neurons (gray), each transparent dot represents a neuron. Vertical and horizontal line plots represent density distributions of each population.

### Outcome-encoding neurons are enriched in frontal cortices

Having identified a distinct subpopulation of RP1 neurons encoding outcome during ongoing movement, we next asked whether these neurons are localized in specific brain regions or uniformly distributed. For each brain region and session, we calculated the fraction of neurons clustered into RP1 and compared it to the overall RP1 fraction within each recording session (18% [17 to 20], mean [95% confidence interval]). RP1 neurons were found in all recorded regions, but their prevalence varied significantly across regions (Fig. 6e). Compared to the overall fraction, RP1 neurons were significantly overrepresented in primary motor cortex (p < 0.001), premotor cortex (p < 0.001), medial prefrontal cortex (p<0.001), mediodorsal thalamus (p < 0.05), sensory thalamus (p < 0.05, LME with animal and session as nested random effect). In contrast, striatum, olfactory areas, and hippocampus contained significantly fewer RP1 neurons than expected by chance (Fig. 6e). Applying the GLM approach to RP1 clusters across brain regions showed that outcome-related activity, predominantly during the reach and before reward consumption, was uniquely encoded in addition to kinematic variables, except in the hippocampus, where this encoding was absent. Finally, we sought to link this outcome-encoding subpopulation to the latent population dynamics and compared the PCA coefficients of RP1 and non-RP1 neurons. RP1 neurons exhibited significantly larger PC1 coefficients than other neurons (p < 0.001 LME, Fig. 6g). These results indicate that the global and local latent population dynamics are strongly shaped by the activity of RP1 neurons, suggesting that the outcome encoding at the latent population level is strongly shaped by the coordinated activity of a distributed subpopulation of outcome-encoding neurons enriched in frontal corticothalamic regions.

## Discussion

Using simultaneous large-scale recordings of neurons across multiple brain regions during a goal-directed reaching task (Fig. 1), we show that movement-related latent dynamics extend beyond classical motor cortical and thalamic regions^23,24,52^. These global dynamics were consistent across individual animals and recording sessions (Fig. 3). We found that both latent population dynamics and a subpopulation of neurons encoded for outcome during reaching, before reward consumption, and that this outcome encoding could not be explained by differences in movement kinematics (Fig. 4-6). Our findings extend prior reports of widespread movement- and outcome-related activity^6,8,33,35,53^ by dissecting a distributed signal that reflects internal outcome expectations during ongoing movement at both single-neuron and population levels.

Low-dimensional latent dynamics during reaching were highly conserved across individuals and shared across cortical and subcortical regions. Previous studies have observed movement-related latent dynamics outside of motor cortex^4,8,27,28^, and that they can be preserved across tasks^54^, time^48^, subjects and shared between motor cortex and striatum^49^. We extend these findings by showing that preserved latent dynamics of goal-directed actions across individuals are not restricted to motor cortex and striatum but can be generalized to other non-motor brain regions as well, such as different thalamic nuclei and the hippocampus. The persistence of this low-dimensional manifold across independently sampled neuronal populations suggests that it reflects a robust and global computational mode that is well suited to integrate sensorimotor and internal signals across brain regions^55^.

A key internal signal during goal-directed movements, such as reaching for a reward, is the continuous change in likelihood of the expected outcome, defined here as the presence or absence of the water droplet. We found a strong modulation of the dominant latent dynamic and of functionally clustered neurons by the presence of reward that existed after controlling for differences in movement kinematics using a GLM (Fig. 4) and a trial-by-trial kinematic similarity analysis (Fig. 5). These results rule out that the outcome encoding is merely a consequence of subtle differences in movement trajectory. Thus, movement-related latent dynamics can be strongly modulated by outcome expectation and are not limited to representing motor preparation or execution^23–25^. This could explain why movement kinematics did not fully explain behavior-related latent dynamics in previous studies^8,23,27,28^.

Most previous work on outcome encoding has focused on reward prediction signals before movement or outcome-related signals during or after reward consumption^8,24,35,37,38,56–58^. Only few studies have analysed outcome encoding during the execution of a goal-directed movement in individual brain regions^59,60^, and to our knowledge, none have done so using simultaneous large-scale recordings across multiple cortical and subcortical areas. We show that anticipated outcome is encoded at the population-level during ongoing movement, and that this signal is widely distributed across cortex, striatum, thalamus, and hypothalamus. The rapid fluctuations of this population activity and its peaking at the onset of reward consumption align with previous findings that the temporal evolution of population dynamics is well suited to represent prospective timing^22,61–63^. Furthermore, our result of action-dependent outcome expectation resembles goal and reward representation during spatial navigation^64–68^ and dopaminergic ramping with reward proximity^30^. These findings support theoretical predictions that internal representations are actionable^39^, meaning that they depend on the execution of an action. For example, the outcome expectation will momentarily change based on the ongoing course of goal-directed action, or in our case the kinematics of a reach. Our study supports action-mediated outcome encoding during forelimb reaching. We propose that this reflects a more general computational mechanism across behaviours, in which the brain integrates sensory feedback and motor state to update internal goal representations in real time.

A functionally defined subset of outcome-encoding neurons contributes to the shared latent dynamics. Using unsupervised clustering and GLM, we identified a subgroup of RP1 neurons showing mixed encoding for outcome and movement that were distributed across many regions. Large-scale multi-region studies have found neurons encoding for various sensorimotor^8,38,69^ or internal variables, such as outcome^8^ , decision^6^ , vigor^70^ , and thirst^12^ , across the brain with many exhibiting mixed selectivity^8,15,38^. However, how this heterogeneous tuning at the single neuron level relates to “multiplexed” global latent subspaces^71^ and the emerging framework of neural manifolds^55^ is not well established, also because simultaneous recording of many brain regions with large neuron populations are still rare. We found that RP1 neurons exhibited high global PC1 coefficients suggesting that they may be key drivers of outcome-dependent latent dynamics across the brain. This would support the hypothesis that the brain-wide manifold emerges from numerous sub-manifolds throughout the brain that by themselves emerge from the activity of distributed and functionally similar neurons^55^.

The enrichment of RP1 neurons in frontal cortex shows that region-specific differences in selectivity and timing exist despite widely distributed encoding^53,69^. We found significantly higher fractions of RP1 neurons in primary motor, premotor and prefrontal cortex compared to other regions, while RP1 neurons were underrepresented in hippocampus, striatum and olfactory regions (Fig. 6). These results are consistent with the established role of motor and prefrontal cortex in outcome encoding^33,72–74^, extending it into the execution phase of goal-directed movements. The high prevalence of outcome-encoding neurons in frontal cortex suggests that these regions could act as key hubs for distributing outcome-related information to functionally coupled neurons across the brain. A similar principle was observed in a multi-region study^8^, where choice signals could be decoded early from sufficiently large numbers of neurons in many areas, but regions with a higher fraction of selective neurons exhibited the earliest and strongest choice information, implying that they originate from those regions and propagate to connected regions. Analogously, we propose that frontal cortical RP1 neurons could generate and disseminate outcome-related population dynamics across the brain. Future experiments that selectively perturb these RP1 neurons will be essential to determine their causal role in driving outcome-encoding latent dynamics in this multi-regional circuit.

Taken together, our findings reveal that outcome expectations are represented during the execution phase of goal-directed movement in a conserved, brain-wide latent dynamic, driven partially by a functional neuronal subpopulation. This work links global neural manifolds with single-neuron mixed selectivity, illustrating how distributed cortical and subcortical networks integrate internal and motor signals in real time. More generally, the observation that outcome representation is continuously broadcasted through a shared latent subspace during movement implies that all regions can process this information based on the principles of reinforcement learning or predictive coding^75–77^. In conclusion, our findings support the emerging perspective that sensory, motor, and cognitive processes are integrated within global neural manifolds that capture dynamic changes of internal representations during ongoing movement^78^.

## Data availability

Upon publication, the data and core analysis code will be made available at https://data.mrc.ox.ac.uk.

## Acknowledgements

We thank Brook Perry, Robert Toth, Ben Micklem, Melissa Serrano, Colin McNamara, Julien Carponcy, Naomi Berry, Shiva Mahdian, Zengcai V Guo, and Jessica Myatt for technical and experimental assistance.

## Author contributions

Conceptualization: YP, AS. Methodology: YP, CL, ST, JS, RS, AL. Formal analysis: YP, ÖO, CL. Investigation: YP, ST. Funding and resources: YP, AS, CS. Data curation: YP, ÖO, CL, ST, JS. Writing – original draft: YP, AS. Writing – review & editing: all authors. Visualization: YP, ÖO, JS. Supervision: AS, CS, AL.

## Methods

### Mice

All experiments were conducted in accordance with institutional guidelines and the UK Animal Scientific Procedures Act (1986) and its associated guidelines. Experiments were performed on eight adult male C57BL/6 mice aged between 3 to 4 months (Charles River). Mice were housed in individually ventilated cages on a 12-hour light/dark cycle. One animal was excluded due to complications during recording sessions, see below.

### Surgery

#### Headplate implantation

Before habituation and training, mice were implanted with a custom-made titanium headplate (Get It Made Ltd, London, 0.7 g) under isoflurane anesthesia (induction 4%, maintenance 1 - 2%). Analgesia was provided using Vetergesic (0.08 mg/kg) administered subcutaneously after induction of anesthesia. The skin on the dorsal surface of the skull was removed. The surrounding skin was secured to the skull using Vetbond (3M, Minnesota, USA). The skull was etched using H_2_O_2_ and a bone scraper to increase adhesion of the headplate that was secured using dental cement (Super Bond C&B, Parkell). A reference screw was wrapped in silver wire and screwed into a craniotomy over the contralateral cerebellum and was fixed in place using a small amount of dental cement (Jet Denture Repair Powder: Lang Dental, Illinois, USA; Meadway Repair Liquid: MR. Dental, Surrey). A custom-designed, 3D-printed shield was glued to the top of the headplate to prevent the animal from reaching recording probes. After implantation, mice recovered for at least a week.

#### Craniotomy surgery

One day prior to recording, craniotomies (∼1.5 mm diameter) were performed under isoflurane anesthesia using a dental bur over the right rostral forelimb area (RFA) (2.5 mm anterior, 1 mm lateral from Bregma), the right caudal forelimb area (CFA) (0.4 mm anterior, 1.3 mm lateral from Bregma) and, in a subset of four animals, at a site directed towards the thalamus (2.5 mm posterior, 1 mm lateral from Bregma). Following surgery, the craniotomies were covered with DuraGel (Cambridge Neurotech) and overlaid with silicone (Body Double, Smooth-On) to form a barrier and protective layer over the exposed brain tissue. During the craniotomy surgery, the hair on the left forelimb was shaved to improve pose estimation of the joints.

### Behavior

#### Reaching task

Before training and recording, mice were habituated to head-fixation over multiple days, beginning with up to 5 minutes of head-fixation and increasing by 5-10 minutes per day. After habituation, mice were trained to perform head-fixed reaches for water droplets (4-5 μl containing 10-15% sucrose) using the left forearm based on a previous study^40^ . Mice had controlled access to water during training and recording, receiving a minimum of 1 ml water per day and enough to maintain a body weight above 85 % of the pre-surgical weight. Training incorporated two components: 1) Mice learned to reach for a droplet upon auditory cue and 2) mice learned to hold the bar for a random interval of 3 to 6 seconds. First, mice were prompted to groom with the left forelimb by application of a small droplet on the left whisker pad which was paired with an auditory cue (3.6 kHz, 80 ms) and a droplet at the spout (4-5 µl). The spout location was initially close to the mouth, allowing mice to lick the droplet. The spout was gradually moved beyond the reach of the tongue, prompting the mice to transition from a grooming motion to a forelimb reaching motion. After mice learned to successfully reach for a water droplet (position: ∼0 mm anterior, ∼5 mm ventral, ∼5 mm lateral to the nose tip), they were trained to hold the metal bar with the left forepaw prior to reaching. Gradually the length of bar-holding time was increased from less than 200 ms to a random interval between 3- and 6-seconds after which the auditory cue and droplet were automatically presented. If mice released the bar prematurely before the cue and droplet presentation, the random interval timer was reset. If mice held the bar until the cue but did not reach or missed the droplet on the reach, the random interval timer was reset after a 10 s timeout. Within 1-2 weeks, mice learned this task and performed over 100 reaches in 30 min. After reaching this threshold, mice underwent craniotomy surgery and subsequent electrophysiological recordings (one recording session per day for 4-7 days).

#### Setup for behavior

We developed a custom-built rig for head-fixed training and recording using Thorlabs parts. The design was inspired by the rig of the International Brain Laboratory (IBL)^79^. The 3D printed headplate holder (Formlabs, Rigid 10k resin) was also adapted from the IBL design. We used the pyControl system for automated control of the task^41^. Specifically, we used the “Lickometer” to electrically detect bar and spout touches. To avoid electrical artifacts during recordings, we integrated a “Comparator Dual Lick Port Detector” (Janelia Experimental Technologies) into the detection circuit. Water droplets (4-5 µl) were dispensed through a gravitation-based system that was controlled by a miniature solenoid valve (Lee Company, LFVA1220210H). Cue tones (3.6 kHz, 80 ms) were delivered via a speaker located at ∼10 cm from the animal. Custom scripts and the pyControl GUI were used to log the behavioral events (bar, spout touch) and automatically trigger the speaker and the solenoid valve.

#### Video recording

Front and side videos were recorded at 100 frames per second using high-speed near infrared enhanced cameras (Ximea MQ013RG-ON or Pixelink PL-D721P). Two LED illuminators (BW 48 LED) were used for infrared lighting. Cameras frames were triggered by the pyControl board at 100 Hz, ensuring synchronization to the pyControl clock. Video data was acquired using StreamPix9 software (Norpix) and encoded in H.264. In one cohort (4 animals), we identified intermittent but systematic frame dropping that caused a consistent drift in the video data relative to the behavioral events. Through *post hoc* comparison of bar release times (measured with pyControl) and paw movement onsets (extracted from Video data using pose estimation described below), we were able to reliably calculate the drift rate and correct for it.

#### Training automated pose estimation model

Using DeepLabCut 2.3.0^42^, we trained one model (ResNet50) per camera (side and front). The side view model was trained on 444 labelled frames and the front view model was trained on 550 labelled frames. The following left forelimb joints were labelled: Shoulder, elbow, wrist, the metacarpophalangeal joint, the proximal interphalangeal joint, and the tip of each digit, excluding the first digit, as it was too small to see. Further labels included the spout, the water droplet, and the proximal interphalangeal joint on the second digit of the right paw. Frames from each animal were included to capture the variance between individual mice. To ensure that the subset of labelled frames reflected most of the positional variance within a session, k-means clustering was applied to the frames in each video and a subset of frames from each cluster was chosen. To further improve generalization and prevent overfitting, the training dataset was also augmented using blurring, flipping, stretching, and partial occlusion of frames. The data was split into training (95%) and test data (5%), and the model trained until the loss plateaued^42,80^. After training, each model was visually inspected and refined through manual labeling of additional frames of underrepresented poses and relabeling of previously labeled frames to increase consistency. The side view model was refined over four training iterations and the front view model was refined over two training iterations. The final side view model had a training root mean squared error (RMSE) of 2.23 pixels and a test RMSE of 2.52 pixels. The last front view model had a training RMSE of 2.18 pixels and a test RMSE of 6.07 pixels. For comparison, human-level performance was previously reported as 2.7 pixels^42,80^.

### 3D triangulation

The trained ResNet50 models were used to label the frames from each video. Each label’s position in 3D space was calculated by direct linear transform (DLT) triangulation with sparse bundle adjustment (SBA) using a wand-based procedure in Argus 3D (Jackson et al. 2016). The intrinsic and extrinsic camera parameters were calibrated using a checkerboard of known spatial dimensions, providing 3D-projections at millimeter scale. Validation through manually labeled frames of checkerboard images yielded a mean scaling offset by -0.017 mm. A 90° corner of the checkerboard was 91.2° in the 3D projection.

#### Video data curation

We visually inspected all triangulated 3D reach trajectories and implemented an automatic filter to exclude mislabeled frames. Estimated coordinates were excluded if they fell in either of these categories: DLC likelihood on front or side camera below 0.5, acceleration exceeding 5 mm/s² (indicates outlier, 95% percentile of physiological acceleration mostly within 1 mm/s²), spatial position outside expected boundaries (in mm: AP [-20 15], DV [-10 15], ML [-10 20]). Linear interpolation was applied to reconstruct excluded coordinates. Further smoothing was applied using a Savitzky-Golay filter with a window size of 5 frames (50 ms). All sessions were visually inspected after automatic curation (Fig. S1).

#### Video-based detection of behavioral times and reach types

All video-based analyses use the proximal interphalangeal joint of the left second digit as the position of the paw, also referred to as the movement trajectory.

#### Reach onset

We used movement trajectory to detect reach onset times, which proved to be more reliable than using the electrical bar release detection, as the comparator occasionally introduced drifts in the detection threshold. We detected the bar holding position as the location with the highest occupancy throughout the session (maximum frequency in AP-ML plane, binned at 0.5 x 0.5 mm). Reach onsets were subsequently detected whenever the paw crossed a threshold of 3 mm anterior to the holding position.

#### Droplet retrieval

We trained the DeepLabCut model for the side view to detect the water droplet, generating a likelihood value for each frame. To detect the time when the animal has successfully retrieved the droplet, we identified instances where the droplet likelihood fell below 0.9 for 3 subsequent time points (20 ms bins) by applying a convolutional kernel to the likelihood timeseries.

#### Goal-directed approaches

To analyse movements irrespective of the trial structure, we detected all movements of the paw towards the droplets throughout the ∼30 min session and defined these as *approaches*. We distinguished them by identifying movements in which the 3D Euclidian distance between the forepaw (proximal interphalangeal joint of the left second digit) and the water droplet decreased. The positions of the joints and water droplets were extracted by the triangulated pose estimation coordinates. To exclude small and noisy movements, we only considered approaches that covered a minimum distance of 2 mm and had an average speed exceeding 0.25 mm/s.

#### Initial reaches

We further distinguished approaches into initial reaches and re-reaches based on the starting position of the paw. Approaches that began within 3 mm of the bar holding position were classified as *initial reaches*. Subsequent re-approaches that do not return to the bar (starting more than 3 mm anterior to the bar) were classified as *re-reaches*. Analysis in this study only focused on the initial reaches.

#### Reach distance

We calculated the reach distance of initial approaches by detecting the maximum distance travelled by an approach within 500 ms after reach onset in the AP dimension. To categorize reach distances like the other properties, we divided them into three groups: 0-5 mm, 5-15 mm, > 15 mm.

#### Droplet consumption (reward)

The end of an approach was determined by the start of the next approach. This represented the time when the paw stopped after moving away from the droplet before the next approach. In a successful approach that included the retrieval of the droplet, this time point represents the moment when the droplet has reached the mouth, which we defined as the start of droplet or reward consumption.

#### Cued

If the approach was performed within 1 second of a cue, it was classified as “cued”. Otherwise, it was “spontaneous”.

#### Outcome

If the droplet was present during the approach, the reach was classified as “rewarded” and the reach period as “reward available”. During absence of a droplet, the reach was “unrewarded”.

#### Grooming

We also detected periods when the animal was performing a grooming movement, instead of a goal-directed reach for a water droplet. These periods could be well-isolated based on the paw moving more dorsal than the typical reach trajectory. When this threshold was crossed within 2 seconds of a reach onset, this approach was classified as a grooming movement.

#### Reach trajectory similarity analysis

In each session, every long reach (>15 mm) performed in the presence of reward (both cued and spontaneous) was paired with all long reaches performed in the absence of reward. We focused on the approach period, defined as the period of continuous decrease in paw-to-droplet distance. For each pair, we computed their differences in speed, trajectory path length, and initial reach angles (measured at the time point when cumulative distance exceeded 10 mm) in both the sagittal (DV–AP) and horizontal (ML–AP) planes. We then calculated the difference in neural activity by subtracting the activity during the unrewarded reach from that of the rewarded reach. Both the activity and kinematic differences of pairs were then averaged across matched time steps. To assess activity differences across different kinematic similarity levels, we grouped the kinematic differences into bins: Δ mean speed: −10 to 10 cm/s (bin size 1 cm/s), Δ trajectory path length: −10 to 10 mm (bin size 1 mm), Δ DV–AP reach angle: −40° to 40° (bin size 10°), Δ ML–AP reach angle: −40° to 40° (bin size 10°). As a control, we repeated the analysis using randomized labels for reward present and absent reaches. Statistical differences in kinematics (e.g. mean speed) between rewarded and unrewarded reaches were assessed using an LME with random intercepts for animal IDs. The model formula was: kinematic_metric ∼ reward + (1 | animalid)

### Electrophysiology

#### Probe preparation

Neuropixels 1.0 probes were sharpened using a repurposed hard drive disk. The ground and reference pads of the probes were shortened and soldered to a wire that connected to the animal and ground. Each probe was labelled with fluorescent DiI (1mg/ml in isopropyl alcohol) before every insertion for *post hoc* probe localization.

#### Neuropixels insertion

We used linear motors (Scientifica IVM) mounted on stereotaxic manipulators (Kopf Instruments 1363A, 3108B) to hold and insert Neuropixels 1.0 probes^81^. Two probes (npx1 and npx2) were held at defined spatial offsets on a custom-printed dual-probe holder (Formlabs, 10k Rigid resin) above the rostral forelimb area (npx 1: 2.5 mm AP, 1 mm ML) and caudal forelimb area (npx2: 1.3 mm AP, 0.4 mm ML) with an DV offset of 0.55 mm. They were inserted simultaneously at 3 µm/s up to a depth of ∼3.5 mm. In animals with thalamic recordings, we inserted a third probe (npx3) to target hippocampus, thalamus, and hypothalamus, inserted at -2.5 mm AP and 1 mm ML with a 14° AP tilt. The speed of insertion was 5 µm/s until ∼4.3 mm depth and then at a speed of 3 µm/s to a total depth of ∼5.3 mm. Once all probes were inserted, they were retracted by 100 µm and allowed to rest in this final position for ∼5 minutes before recording. After each recording session, probes were cleaned using Terg-a-zyme (1%, Sigma Aldrich), isopropyl alcohol and dH_2_O.

#### Data acquisition

Electrophysiological data were acquired using OpenEphys software in binary format. Default settings were used for gain (AP band 500x, LFP band 250x), sampling rate (AP band 30 kHz, LFP band 2.5 kHz), and channel mapping (384 electrodes from tip). Recordings were referenced to the tip of the probe. Data were synced using random TTL pulses generated by pyControl. The Rsync module of pyControl was used to synchronize the electrophysiological data, behavioral events and the triggered video frames for downstream analyses.

#### Histology and imaging

After recordings, mice were anesthetized with a terminal intraperitoneal injection of pentobarbital (200 mg/kg). Each mouse was transcardially perfused with 0.1 M phosphate buffered saline (PBS), followed by 4 % paraformaldehyde solution (PFA) in PB. The brains were removed, post-fixed for 2 hours in 4 % PFA in PB at room temperature, then washed with and stored in 0.1 M PB overnight at 4 ° in the dark. The following day, brains were transferred into 10 % sucrose in 0.1 M PB for storage at 4 ° overnight and finally transferred to 30 % sucrose in 0.1 M PB until sectioning. Brains were sectioned coronally at 50 µm per slice using a freezing microtome (Epredia HM 450) and stored in PB-Azide until mounting for imaging. Brightfield and DiI images were acquired using the Cy3 imaging protocol (Zen Blue software, excitation using 532 nm laser line) on an epifluorescent microscope (Zeiss Axio Imager M2) and a 5x objective.

#### Region assignment

Probe tracks were reconstructed using the SHARP-Track toolbox^82^. Selected images were processed and aligned to the matching coordinates of the Allen mouse brain reference atlas. Probe tracks were then manually traced based on fluorescent tracks. Different probe trajectories were assigned to recording sessions based on matching of the reconstructed and the experimentally documented (manipulator coordinates) entry point into the brain. In cases of fewer recovered probe tracks than experimentally performed insertions, the most proximal trajectory was assigned. Adapted SHARP-Track code was used to map Allen Mouse Brain Common Coordinate Framework (Allen CCF) region labels to channels along the length of the Neuropixels probe. For the initial assignment, the “probe_length” parameter was based on the final coordinates of the linear motor. No scaling factor was applied.

To further align the region boundaries based on electrophysiological landmarks along the Neuropixels probe^83^, we designed a custom GUI to visualize relevant spike- or LFP-related data along the length of the probe: Spike raster, firing rate, cell count, waveform peak amplitude, AP band activity (number of crossing below -100 µV in 30 s interval), raw LFP (after subtracting median across channels), LFP power (Welch’s power spectral density estimate), LFP power normalized across channels, LFP power relative change across channels, LFP cross correlation across channels (see Fig. S3e for subset of those parameters). We visually inspected each probe insertion and shifted the probe to align region boundaries to characteristic electrophysiological landmarks as reported previously^83^: On Neuropixel-1 (npx-1: premotor cortex, medial prefrontal cortex, orbitofrontal cortex, olfactory areas), we aligned the boundary between cortex and olfactory areas to the sharp and consistent increase in low-frequency LFP power (Fig. S3e). On npx-2 (primary motor cortex and dorsal striatum), we aligned the boundary between white matter and caudaputamen (CP) to the sharp and consistent increase of high frequency LFP power (Fig. S3e). Here, the corpus callosum and associated fibers are often demarked by a drop in cell count and overall LFP power. On npx-3, we aligned the boundary between hippocampus and thalamus to the sudden disappearance of within-hippocampal cross-correlation (molecular layer above and below pyramidal layer of dentate gyrus). This shift also aligned the “cross” of the LFP cross correlation to the center of the dentate gyrus pyramidal layer and corresponded to an increased spike amplitude in thalamic nuclei.

Overall, we found that a shifting of the region boundaries was sufficient so that other region boundaries also showed consistent and aligned changes in different parameters. To account for the expected imprecision, we grouped CCF region labels into broader region groups. White matter regions and channels outside the brain were assigned as “Other” and excluded from analysis (only visualized in Fig. 2c).

#### Session curation

We initially recorded 42 sessions from 8 mice (Fig. S4a). After visual inspection of all data, we excluded four sessions due to the following technical reasons: One session was missing video files, one session had faulty bar hold detection, one animal only had two recording sessions, as we had to abort the recordings due to experimental complications. Due to the low number of sessions and incomplete data in the second recording session, we excluded this animal. The remaining data (38 recording sessions from 7 mice, 4-7 sessions per animal) were used throughout all analyses.

#### Spike sorting and quality control

Automated spike sorting was carried out using KiloSort 3.0^84^. Using default parameters on the included 38 recording sessions of ∼30 min duration each, we obtained a total of 76,163 isolated clusters of which 44,479 were labeled as “good” by Kilosort (51%, Fig. S4a). Quality metrics were automatically calculated for all isolated clusters using the toolbox Bombcell^85^ with default parameters, except that the minimum number of spikes was set at 50. For all analyses, we only included units that were labelled “good” by both Kilosort and by Bombcell which yielded a total of 22,036 (29% of all isolated clusters). To validate the spike sorting and the automated quality control, we visually inspected 2,073 units (∼9% of all units, at least 20 per probe insertion) using phy (cite) and found that 94% ± 0.8% (mean across session ± SEM) could be confirmed as a “good” unit.

#### Data integration and pipeline code

All data were integrated into the CellExplorer^86^ data framework for further analysis. The Rsync module by pyControl was used to synchronize video data, behavioral data, and spike times, using the time of the electrophysiological data as reference. A custom MATLAB-based data analysis pipeline was developed for data integration, processing, analysis, and visualization. Data were visualized using the gramm toolbox^87^ or custom-written code. Large language models were used to assist coding, mainly to refine human-generated code. AI-generated code was checked by the authors and validated through control analyses.

#### Modulation index

The modulation index was calculated by the CellExplorer processing module^86^. It represents the relative increase in trial-averaged mean firing rate within 500 ms after the cue relative to the mean baseline firing rate within 250 ms before the cue. The p-value was calculated using a two-sample Kolmogorov-Smirnov test between the firing rate distribution and the post-cue and baseline period across trials. Significantly tuned neurons were identified after Bonferroni correction for multiple comparisons (alpha = 0.05).

#### Principal component analysis across reach types

##### Data preprocessing and reach types

Firing rates of each neuron were binned at 20 ms and smoothed using a Gaussian kernel (σ = 20 ms). Each neuron’s firing rate was z-score normalized across the full recording session to remove differences in overall firing rates. We obtained qualitatively similar results in a control analysis without any normalization (Fig. S4f). Neural activity was segmented into trial-aligned epochs from −1.5 to +3 s relative to movement onset. Trials were categorized into five reach types based on cue, reach distance, and outcome as defined above: cued long rewarded reaches, spontaneous long rewarded reaches, spontaneous long unrewarded reaches, spontaneous short unrewarded reaches, and grooming. Trials were excluded if the analysis window extended beyond session boundaries and conditions with fewer than five trials within a session were not analyzed.

##### PCA within-reach type with cross-validation

To assess the reliability of latent dynamics, trials were randomly split into 80% training and 20% test sets. Trial-averaged activity was computed separately for each split, and PCA was performed on the training average. Both training and test trial-averaged population trajectories were projected into the PCA subspace defined by the training data. For each principal component, the resulting time courses from training and test data were then compared by computing their correlation, providing a measure of how reliably each component was reproduced in held-out data. Similarity between trajectories was quantified for each principal component using the squared Pearson correlation coefficient (R²).

##### PCA projection across reach types

To compare latent dynamics across reach types, PCA was computed on the trial-averaged activity of a reference reach type (“training subspace”), and the activity from other reach types was projected into the same subspace (“projected latent dynamics”). Similarity between the reference and projected trajectories was quantified per component using R². This approach assessed the extent to which population dynamics generalize across reach types.

##### Alignment and within-session scaling

Because the sign of principal components is arbitrary, component signs were aligned across sessions based on the cued reach. To enable comparison of latent amplitudes across sessions and conditions, projected latent dynamics were scaled within each session by the peak absolute amplitude of the corresponding component in the training PCA subspace, computed within 500 ms post-movement time window. This normalization preserved temporal structure while allowing comparison of relative amplitudes across reach types.

#### Across-session similarity using CCA and Procrustes distance

To compare the similarity of latent dynamics across sessions, we performed separate PCA for each reach type within each session and retained the first 10 principal components. For each reach type, pairwise comparisons were then computed between all session pairs using canonical correlation analysis (CCA) and Procrustes analysis. CCA was used to quantify the correlation between aligned latent trajectories across sessions, and the canonical correlation coefficients were extracted for the first 10 components. Procrustes distance was used to quantify the geometric similarity of latent trajectories across sessions independent of translation, rotation, and scaling. Session pairs were classified as within-animal or across-animal based on the identity of the recorded mouse. As a control, we repeated the same across-session CCA and Procrustes analyses using latent trajectories derived from PCA on randomly sampled 4.5 s segments of neural activity from each session. For each session, 100 random control trajectories were generated, pairwise similarity was computed across sessions, and the resulting CCA coefficients and Procrustes distances were averaged across shuffles. This control provided an estimate of the similarity expected from task-independent population covariance.

#### Region-specific principal component analysis

To characterize region-specific population dynamics, we performed PCA separately for each brain region using trial-averaged activity pooled across sessions. Firing rates of each neuron were binned at 20 ms, smoothed using a Gaussian kernel (σ = 20 ms), and z-score normalized across the full recording session, such that each neuron had zero mean and unit standard deviation across time. Neural activity was aligned from −1.5 to +3 s around movement onset. For each session and brain region, trial-averaged responses were computed separately for each reach type, requiring at least five valid trials per condition. These session-level averages were then concatenated across sessions within each region to obtain a pooled activity matrix of time bins × neurons, and PCA was applied to this matrix. To compare latent dynamics across regions and reach types, the sign of each principal component was aligned within each reach type by maximizing its correlation with the mean trajectory of the corresponding component across all other regions. For the cross reach type analysis, PCA was trained separately within each brain region on pooled activity from cued long rewarded reaches. Trial-averaged activity from the other reach types was then projected into this same region-specific cued-long PCA subspace on a session-by-session basis using the corresponding region-specific neurons.

#### Generalized linear models

We fitted Generalized Linear Models (GLMs) to explore how task variables were encoded in different types of neural activity. The model structure was based on Engelhard et al. (2019)^50^ . As the dependent variable, we used the continuous expression of neural activity. To obtain the time-varying expression of latent PCA dynamics, we projected each neuron’s continuous and normalized firing rate timeseries onto the previously computed PC/dPC weights by multiplication. We then performed multiple linear regression against a set of behavioural predictors. GLMs were trained on trial-wise data and evaluated with 5-fold cross-validation. A trial covered the period between the reach onset and offset and included up to one second of baseline activity before and after a reach.

We used two classes of predictors: 1) continuous predictors: distance of the second digit to the bar, and its speed; 2) event predictors: movement-onset, cue, reward-presence, and reward-consumption. Following Engelhard et al. (2019), event predictors were convolved with spline basis sets generated using the bs package in R (200 ms long splines with 7 degrees of freedom for movement-onset and cue, 1 sec long for the reward-consumption predictor). Specific spline subsets were assigned as follows: movement-onset (splines 2–6), cue (4–6), and reward-consumption (1–5). Finally, reward presence predictor was a step function set to 1 when droplet likelihood exceeded 0.9. All other model parameters followed Engelhard et al. (2019). Predictor contributions were estimated using the “no-refit” method, in which each predictor was removed from the full model by zeroing its regression coefficient. The unique explained variance (R²) for each predictor was computed as its relative contribution multiplied by the full model’s R². Predicted activities were calculated using the full model.

##### Collinearity analysis

To assess multicollinearity among predictors, we computed variance inflation factors (VIF) and pairwise correlations on the trial-epoch design matrix (bar distance, speed, reward presence, reward consumption, cue, and bar-off). VIFs were computed by regressing each predictor on all others and applying 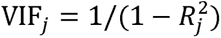.

#### Subpopulation clustering analysis

To identify functional clusters within the neural activity in an unsupervised manner, we applied the Rastermap algorithm (Stringer et al., 2025) to session-wide normalized firing rates all recorded neurons. We set the locality parameter to 0 to enable sorting without region separation, used a time lag window of 10, and tested different values for the cluster number parameter (n_cluster = [4:1:8]). While different cluster numbers resulted in comparable results, we proceeded with n_cluster = 6 as it maximized the variance explained by the full GLM model and the highest unique contribution of reward-presence. For each session, the cluster with the highest R² value explained by the reward-presence predictor were labelled as the Reward Presence 1 (RP1) cluster.

##### Region enrichtment

To assess whether RP1 units, corresponding to the cluster with the highest reward-presence R², were enriched in specific brain regions, for each region we calculated the fraction of units labelled as RP1. This was compared to the RP1 fraction per session. To test for statistical RP1 enrichment across regions, we fit an LME with animal ID and session as nested random effect. The model formula was: RP1% ∼ region + (1 | animalid) + (1 | animalid:session)

