## Supplementary figures 1-5 for "Distributed encoding of action-mediated outcome drives consistent population dynamics during goal-directed reaching"

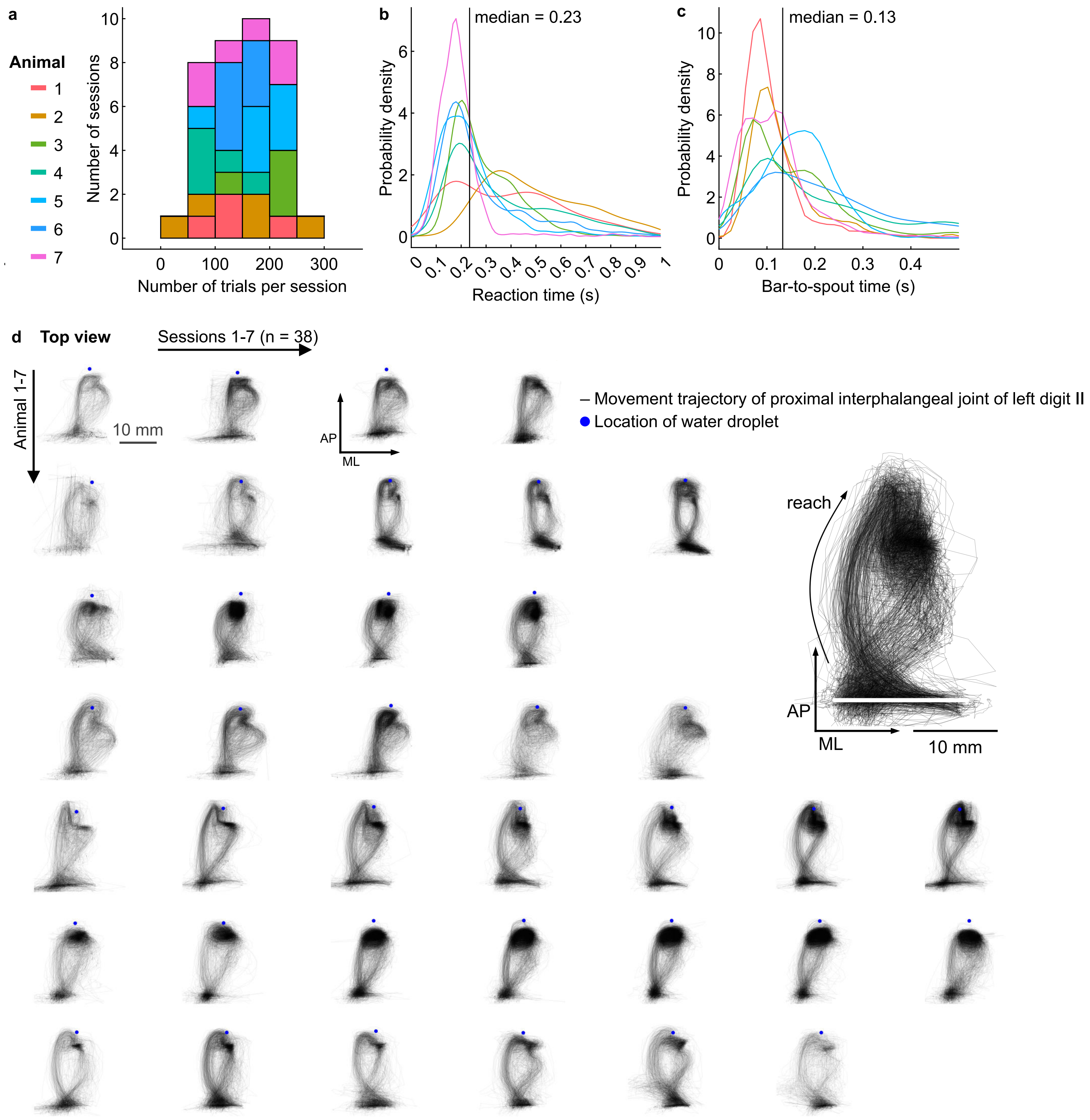


**Supplementary figure 1: Task performance and movement trajectories. (a)** Stacked histogram shows distribution of number of correctly performed cued reach trials per 30-minute recording session, across sessions and animals (color). **(b)** Probability density plot shows distribution of reaction time across trials, pooled within each animal (colored lines). Vertical line indicates median of 0.23 s. **(c)** Same plot as in panel b, showing distribution of movement time from bar to first spout touch.

**(d)** Top view of movement trajectory of left digit II during full 30-minute recording session (transparent traces) after data curation (see methods). Individual sessions (columns) of different animals (rows) are shown. Blue dot indicates location of water droplet. Location of bar is at the bottom. Note the consistent movement trajectories within animals. The darker frequently traversed space close to the droplet arise from multiple re-reaches between mouth and spout. Top right shows enlarged example from one session.

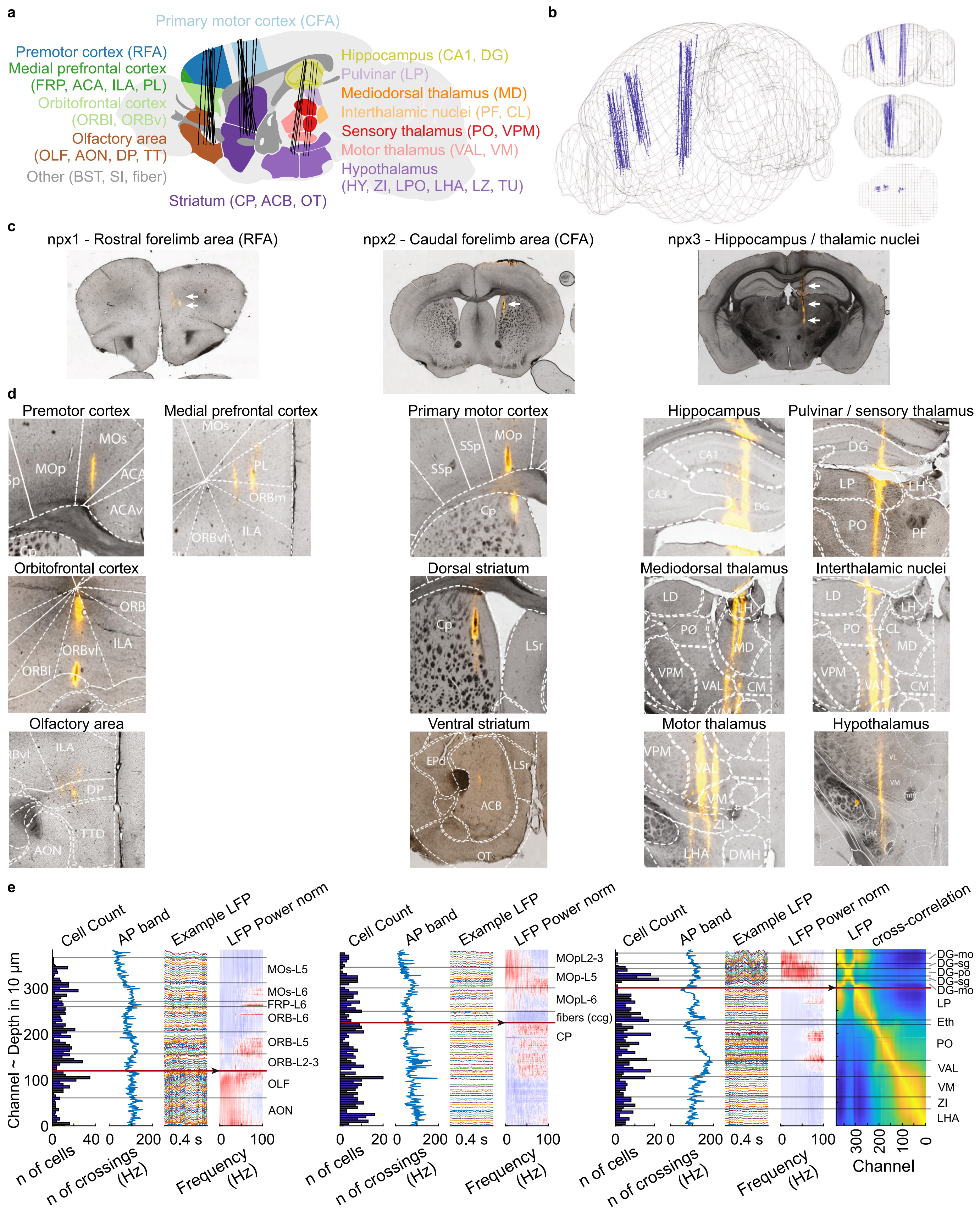


**Supplementary figure 2: Probe localization based on histology and electrophyisology. (a)** Same panel as in Figure 1, shown for reference: Schematic of sagittal brain atlas showing recorded region categories in color. Abbreviation in parentheses indicate subregions according to Allen Mouse Common Coordinate Framework (CCF). Black lines represent Neuropixels trajectories from different sessions approximating positions of active electrodes. **(b)** Schematic shows probe trajectories (blue lines) in normalized 3D brain model (gray wireframe) based on labelled points (blue dots). Probe tracts localized and plotted using the SharpTrack toolbox. Right side shows same probe tracts on saggital , frontal, and horizontal plane. **(c)** Brightfield imaging of horizontal brain slices showing fluorescent DiI probe tracts from each of the three insertion sites (orange, white arrows). **(d)** Example brightfield images showing the probe tracts (orange) going through each broader brain region. Overlaid and registered atlas boundaries and region labels are shown in white. **(e)** Plots show example data from individual sessions from the three insertion sites mapped across the laminar profile of the probe (channels 1-384, probe tip at 0). Cell count: Histogram of number of cells recorded on each channel (bin size: 5 channels). AP band: Line plot shows frequency of AP band trace crossing -100 µV threshold. Example LFP: Colored traces represent 400 ms of LFP trace from every 5[th] channel. LFP Power norm: Heatmap shows power spectrum densities, normalized across channels. The sharp increase in relative power are used as electrophysiology landmarks for RFA and CFA probes (black arrow). LFP cross-correlation: Heatmap shows cross-correlation of LFP traces between each channel. Note the strong cross-correlation between the two granule cell layers in the dentate gyrus (DG-sg). The bottom of this „cross" was used as landmark for the thalamic probe.

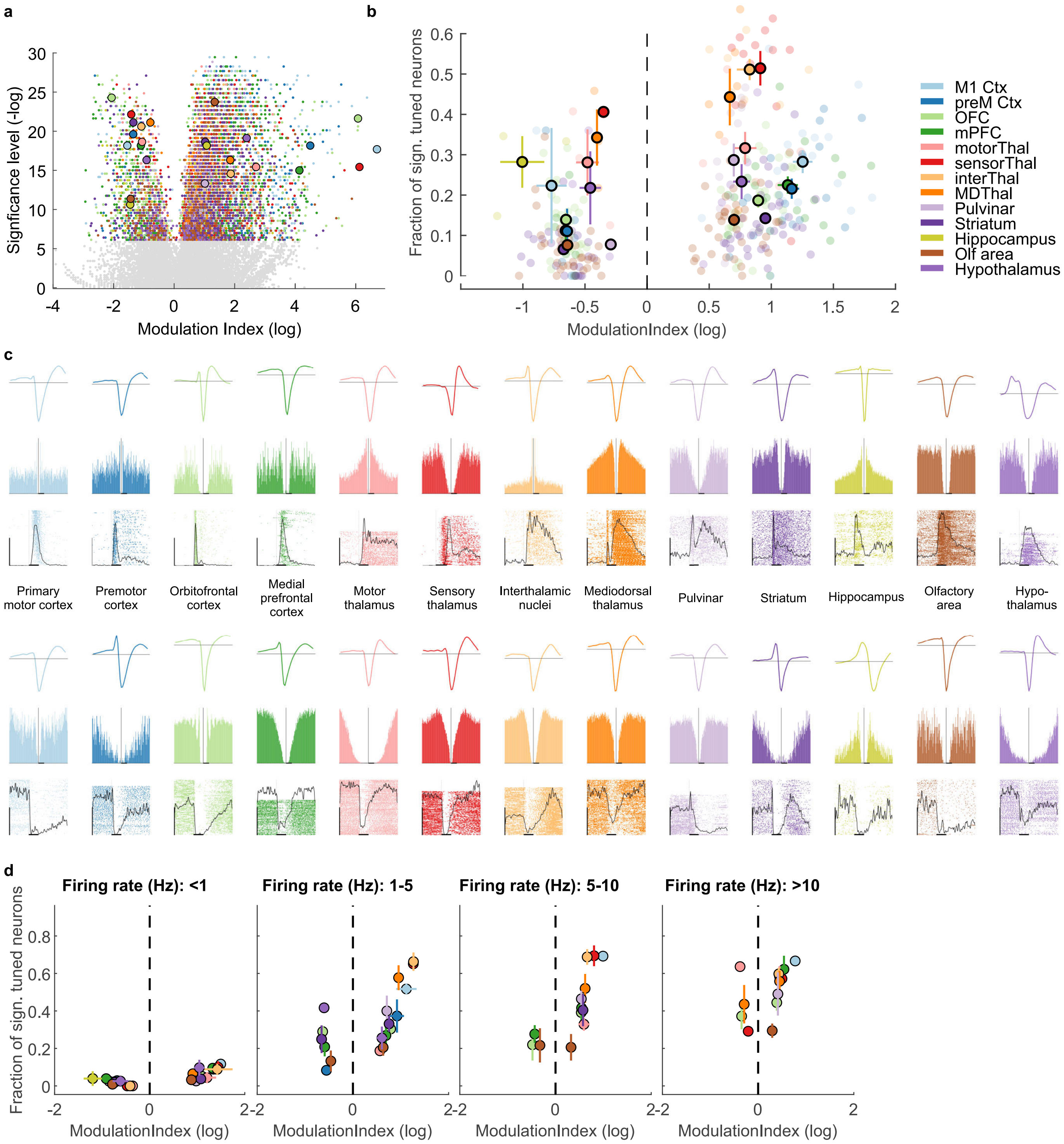


**Supplementary figure 3: Example spike rasters across brain regions. (a)** Same panel as in Figure 1 with more highlighted example neurons. Volcano plot shows cue modulation index (log) and significance level (-log10) for each recorded neuron, color-coded by its brain region. Grey neurons are not significant after Bonferroni correction ($p > 1.1 \times 10^{-06}$). Circled dots indicate example neurons from each region shown in panel c and d (matching colors). **(b)** Scatter plot shows fraction of significantly positively modulated (index > 1) and negatively modulated (index < 1) neurons in each region (colour). Each transparent dot indicates an individual recording session, circled dots represent mean and standard error of mean across sessions per region. **(c)** Spike template waveforms (top row), auto-correlograms (middle row, horizontal scale bar: 10 ms) and spike rasters (bottom row, aligned to time of cue, vertical scale bar: 100 trials, horizontal scale bar: 500 ms) shown for example units (each column) with positive modulation index that were highlighted in panel a. Color code represents the brain region. Black line in spike raster represents scaled average firing rate across trials. (d) Same plot as in panel b, showing region-wise averages of fraction of significantly tuned neurons and modulation index, averaged within different firing rate bins.

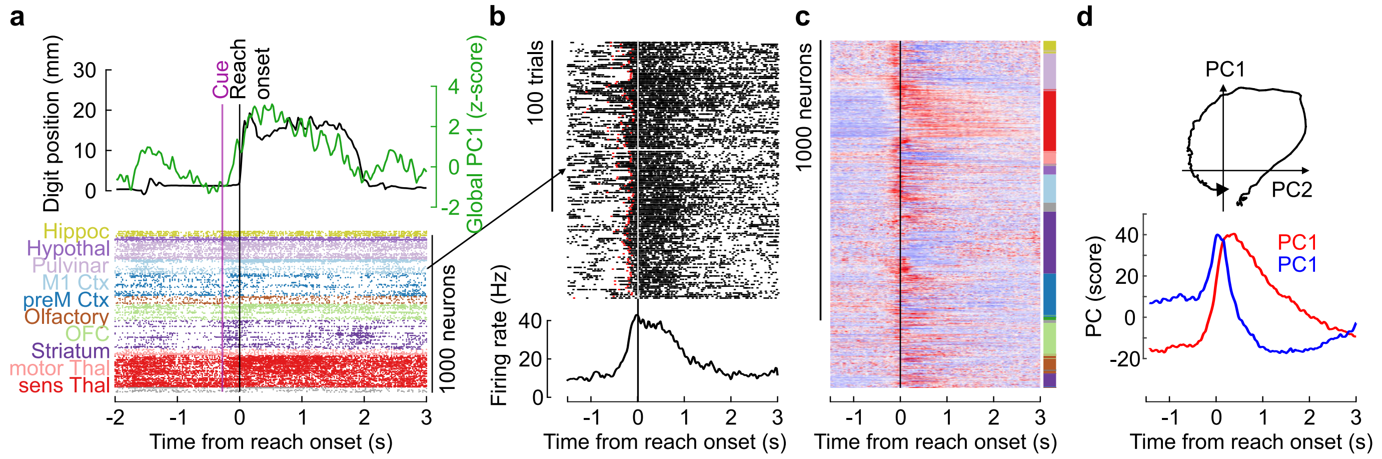


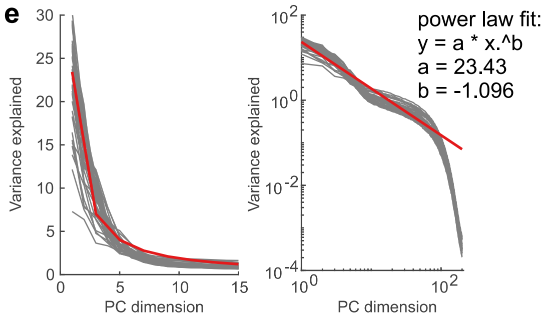


**Supplementary figure 4: Principal component analysis and consistency across movement types.** **(a)** Single-trial example showing change in digit position (black) and PC1 latent dynamic (green) around behavioural events (cue, reach onset, spout touch). Raster plot below shows spiking activity of all simultaneously recorded neurons, color coded by brain region. **(b)** Spike raster plot of an example neuron across trials (rows) aligned to time of reach onset. Red dots indicate relative time of preceding cue. Trial-averaged firing rate is shown below. **(c)** Peri-event time histogram shows z-scored firing rate (color code) of all neurons (rows) in example session. Colored bar on right indicates region. **(d)** Temporal dynamics of principal components 1 and 2 of example session, computed on trial-averaged population activity. **(e)** Scree plot showing variance explained across PC1-15, gray lines represent individual sessions. Red line represents power law fit. Right panel shows variance explained across PC dimensions in a log-log plot. Note that the decay is well approximated by a power law decay (red line).

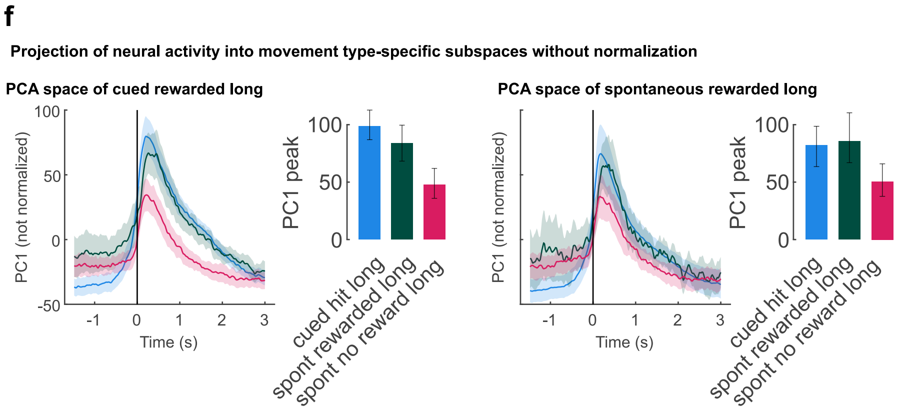


**(f)** Trial-averaged PC1 traces of different reach types (colors) were obtained by projecting activity into the PCA subspace trained on either rewarded cued long reach (left) or rewarded spontaneous long reach (right) without any normalization. Thick lines represent across-session mean. As in panel d, within-type projections show 20/80 cross-validation. Bar plots show average of normalized PC1 peak amplitude across sessions, error bars indicate 95% CI.

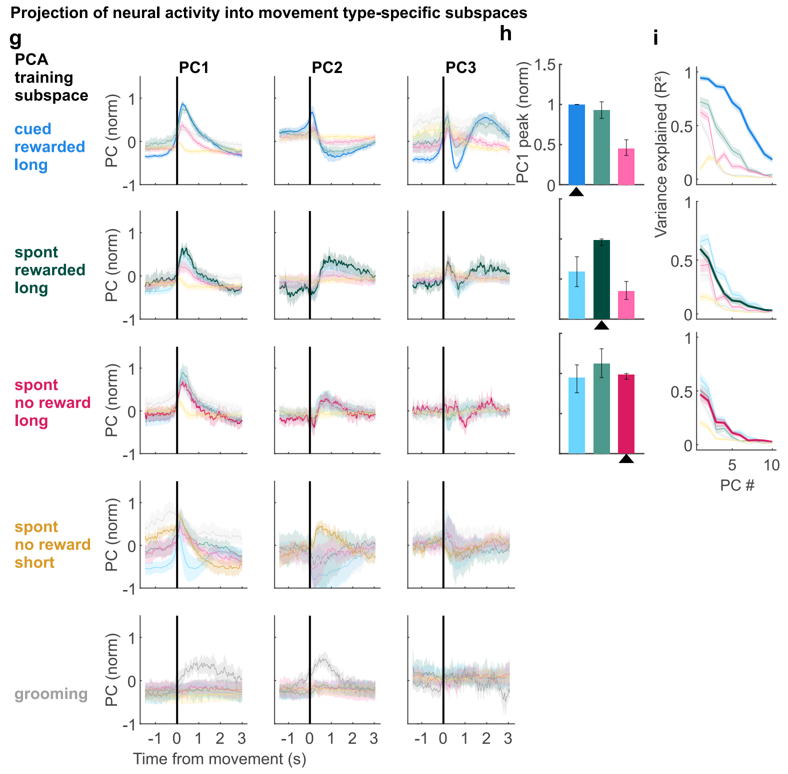


**(g)** Principal component (PC) trajectories of the top three components (columns) extracted from neural population activity aligned to movement onset. Each row shows PCs trained on trial-averaged activity from a specific movement type (indicated on the left), and then projected into activity from multiple movement types (color-coded within each row). Thick colored lines represent activity from the same movement type used for training the PCA (within-type projections), while thinner lighter lines represent cross-type projections. Traces show the across-session average; shaded regions indicate bootstrapped 95% confidence intervals. Black vertical lines mark movement onset (t = 0 s). **(h)** Bar plots show the mean peak amplitude (across sessions) of the first principal component (PC1) within 1 s of movement onset for three long reach types (colors), projected into PCA subspaces trained separately on each of these movement types indicated by black triangles in each row. Light colors represent cross-type projection, dark colors within-type projection. Error bars indicate bootstrapped 95% confidence intervals. Statistical comparisons were performed using Wilcoxon signed-rank tests (* $p < 0.05$, *** $p < 0.001$). **(i)** For the PCA trained on cued long reaches line plots show the mean fraction of variance ($R^2$) of the top ten PCs across sessions. $R^2$ represents how much variance of the trial-averaged neural activity of different movement types (colors) is explained when projected into this PCA subspace (cross-type projection = thin lines). Thick lines indicate within-type projections, where $R^2$ quantifies the variance explained by each principal component when projecting activity of held-out test trials (20%) into the PCA subspace calculated from training trials (80%) of cued long reaches.

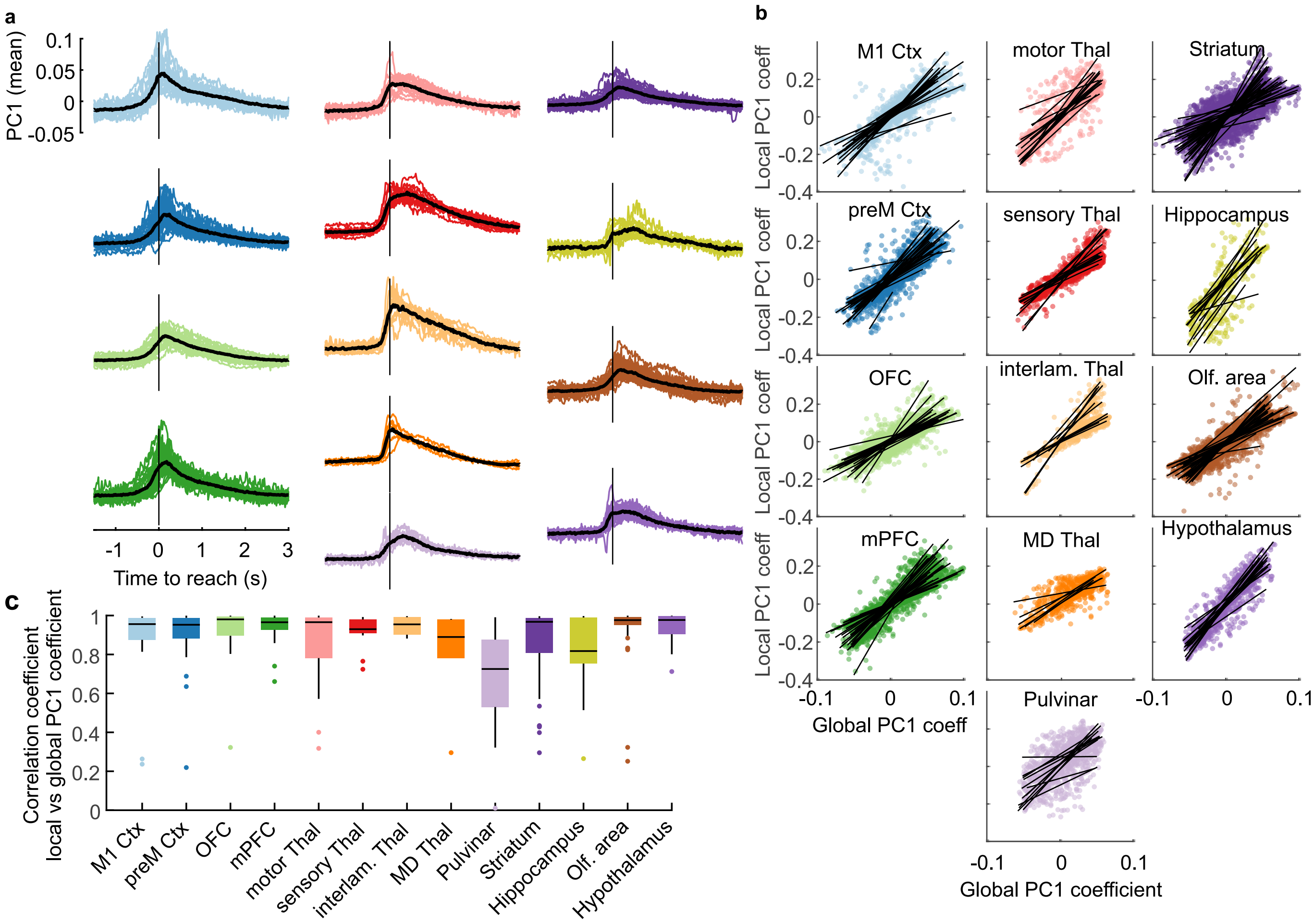


**Supplementary figure 5: Correlation of global and local PCA coefficients.** (a) Region-specific expression of PC1 dynamic, individual lines represent different sessions, colors correspond to region annotation in (c). Black line represents mean across session. Vertical line indicates time of reach onset. **(b)** Scatter plots show the loading weights(coefficients) of each neuron (dots) for the PC1 dimension, calculated either by PCA of all neurons within the recording (global, x-axis) or only by PCA of only neurons of the same region (local, y-axis). Different colors and plots represent different brain regions (bold label to the right). Black lines represent linear fits of individual recording sessions. **(c)** Box plots show the correlation coefficient calculated from the PC1 coefficient of PCA, either from all neurons or from neurons in a specific brain region (x-axis and color code). Each data point represents a single recording session.
